# Transcripts’ evolutionary history and structural dynamics give mechanistic insights into the functional diversity of the JNK family

**DOI:** 10.1101/119891

**Authors:** Adel Ait-hamlat, Diego Javier Zea, Antoine Labeeuw, Lélia Polit, Hugues Richard, Elodie Laine

## Abstract

Alternative splicing and alternative initiation/termination transcription sites, have the potential to greatly expand the proteome in eukaryotes by producing several transcript isoforms from the same gene. Although these mechanisms are well described at the genomic level, little is known about their contribution to protein evolution and their impact at the protein structure level. Here, we address both issues by reconstructing the evolutionary history of transcripts and by modeling the tertiary structures of the corresponding protein isoforms. We reconstruct phylogenetic forests relating 60 transcripts from the c-Jun N-terminal kinase (JNK) family observed in 7 species. We identify two alternative splicing events of ancient origin and show that they induce subtle changes on the protein’s structural dynamics. We highlight a previously uncharacterized transcript whose predicted structure seems stable in solution. We further demonstrate that orphan transcripts, for which no phylogeny could be reconstructed, display peculiar sequence and structural properties. Our approach is implemented in PhyloSofS (Phylogenies of Splicing Isoforms Structures), a fully automated computational tool freely available at https://github.com/PhyloSofS-Team/PhyloSofS.

## Introduction

Alternative splicing (AS) of pre-mRNA transcripts and alternative transcription initiation/termination are essential eukaryotic regulatory processes. They can impact the regulation of gene expression, for instance by introducing changes in the three prime untranslated region [1]. Or they can directly modify the content of the coding sequence (CDS) [2], leading to different protein isoforms. Virtually all multi-exons genes in vertebrates are subject to AS [3] and about 25% of the AS events (ASEs) common to human and mouse are conserved in other vertebrates [6, 7, 8]. This suggests an important role for AS in expanding the protein repertoire through evolution. AS has also gained interest for medicinal purpose, as the ratio of alternatively spliced isoforms is imbalanced in several cancers [4, 5].

The extent to which the ASEs detected at the gene level actually result in functional protein isoforms in the cell remains largely unknown. Transcriptomics and proteomics studies suggested that most highly expressed human genes have only one single dominant isoform [9, 10], but the detection rate of these experiments is very difficult to assess [11] and likely suffer from strong experimental detection bias [12]. A recent analysis of ribosome profiling data suggested that a major fraction of splice variants is translated, with direct implications on specific cellular functions [13]. Moreover, a large scale assessment of isoforms present in the cell revealed that the majority of isoform pairs share less than 50% of their interactions [14]. From a structural perspective, very few alternatively spliced isoforms have been characterized and are available in the Protein Data Bank (PDB) [17, 18]. In human, it was shown that the boundaries of single constitutive exons or of co-occurring exon pairs tend to overlap those of compact structural units, called protein units [19]. While tissue-specific alternatively spliced exons tend to be enriched in disordered regions containing binding motifs [20], it was suggested that splicing events may induce major fold changes [21, 22]. Along this line, a few cases of isoforms displaying domain atrophy while retaining some activity have been reported [23].

The elusiveness of the significance of AS for protein function and fold diversification though evolution has stimulated the development of knowledge bases, such as APPRIS [24] and Exon Ontology [25]. They provide functional and sequence-based information at the level of the transcript or the exon. A method reconstructing transcripts’ phylogenies was also proposed and proved useful for enhancing transcriptome reconstruction from ESTs and investigating proteins functional features (domains, sites) [15, 16].

In this work, we combine sequence- and structure-based information to shed led on the evolution of AS. We have developed PhyloSofS (Phylogenies of Splicing Isoforms Structures), an automated tool that infers plausible evolutionary scenarios explaining an ensemble of transcripts observed in a set of species and predicts the tertiary structures of the protein isoforms. We show how PhyloSofS can be used to provide insight on the contribution of AS to the evolution of the c-Jun N-terminal kinase (JNK) family and on the molecular mechanisms underlying AS functional outcome. JNKs are essential regulators that target specific transcription factors (c-Jun, ATF2...) in response to cellular stimuli. The deregulation of their activity is associated with various diseases (cancer, inflammatory diseases, neuronal disorder...) which makes them important therapeutic targets [29]. About ten JNK splicing isoforms have been documented in the literature [30]. They were shown to perform different context-specific tasks [31, 32, 33, 34] and to have different affinities for their substrates [35, 36]. By reconstructing the phylogeny of JNK transcripts across seven species, we identify two ASEs of ancient origin. We further identify key residues that may be responsible for the selective recognition of JNK substrates by different isoforms and characterize the behaviour of these isoforms in solution by biomolecular simulations. One of the ASEs involves a 80-residue deletion and has never been documented before. We find that its predicted structure is stable in solution. Both ASEs are supported by sequencing evidence from transcriptomics studies.

Our work allows to put together, for the first time, two types of information, one coming from the reconstructed phylogeny of transcripts and the other from the structural modeling of the produced isoforms, and this to shed light on the molecular mechanisms underlying the evolution of protein function. It goes beyond simple conservation analysis, by dating the appearance of ASEs in evolution, and beyond general structural considerations regarding AS, by characterizing in details the isoforms’ shapes and motions. We find that the effect of functional ASEs on the structural dynamics of the isoforms may be subtle and require such a detailed investigation. Our results also open the way to the identification and characterization of new isoforms that may be targeted in the future for medicinal purpose.

### New approaches

PhyloSofS can be applied to single genes or to gene families. Given a gene tree and the observed transcripts at the leaves (Fig. 1a, on the left), it reconstructs a phylogenetic forest embedded in the gene tree (Fig. 1a, on the right) representing plausible evolutionary scenarios explaining the transcripts. Each tree in the forest (in orange, green or purple) represents the phylogeny of one transcript. In other words, each root indicates the appearance of a new transcript and its corresponding ASE(s). Transcript losses are possible (triangles in Fig. 1a), and the exon usage of a transcript can change along the branches upon the inclusion or exclusion of one or several exons (which we refer to as “mutations”). The underlying evolutionary model is comprised of two levels, following [15]. At the level of the gene, exons can be absent, constitutive, or alternative (*i.e.* involved in at least one ASE), whereas at the level of the transcript, exons are either present or absent. The cost associated to the mutation of an exon naturally depends on its impact on the status of the exon at the gene level. For instance when the gain of an exon at the gene level shifts its status from absent to constitutive, the mutation will not be penalized (see *Methods* and Table 2). PhyloSofS algorithm seeks to determine the scenario with the smallest number of evolutionary events, following the maximum parsimony principle. It is inspired from that reported in [15]. Our main contribution was to develop heuristics in order to treat complex cases in a computationally tractable way. Specifically, we have implemented a multi-start iterative strategy combined with a systematic local exploration around the best current solution to efficiently search the space of phylogenetic forests (see *Methods* and S1 Fig). Moreover, we have designed a branch-and-bound algorithm adapted to the problem of assigning transcripts between parent and child nodes (see *Methods* and S1 Text). The reconstructed forests are provided with a user-friendly visualization (Fig. 1b-c). In addition to phylogenetic reconstruction, PhyloSofS predicts the 3D structures of the protein isoforms. The predictions are performed based on comparative modeling using the HH-suite [37]. Furthermore, PhyloSofS annotates the generated models with sequence (exon boundaries) and structure (secondary structure, solvent accessibility, model quality) information. For example, it is very easy to visualize the location of each exon on the modeled structure. Here, we present the application of PhyloSofS to the c-Jun N-terminal kinase (JNK) family across 7 species (*H. sapiens*, *M. musculus*, *X. tropicalis*, *T. rubripes*, *D. rerio*, *D. melanogaster* and *C. elegans*).

**Figure 1.**
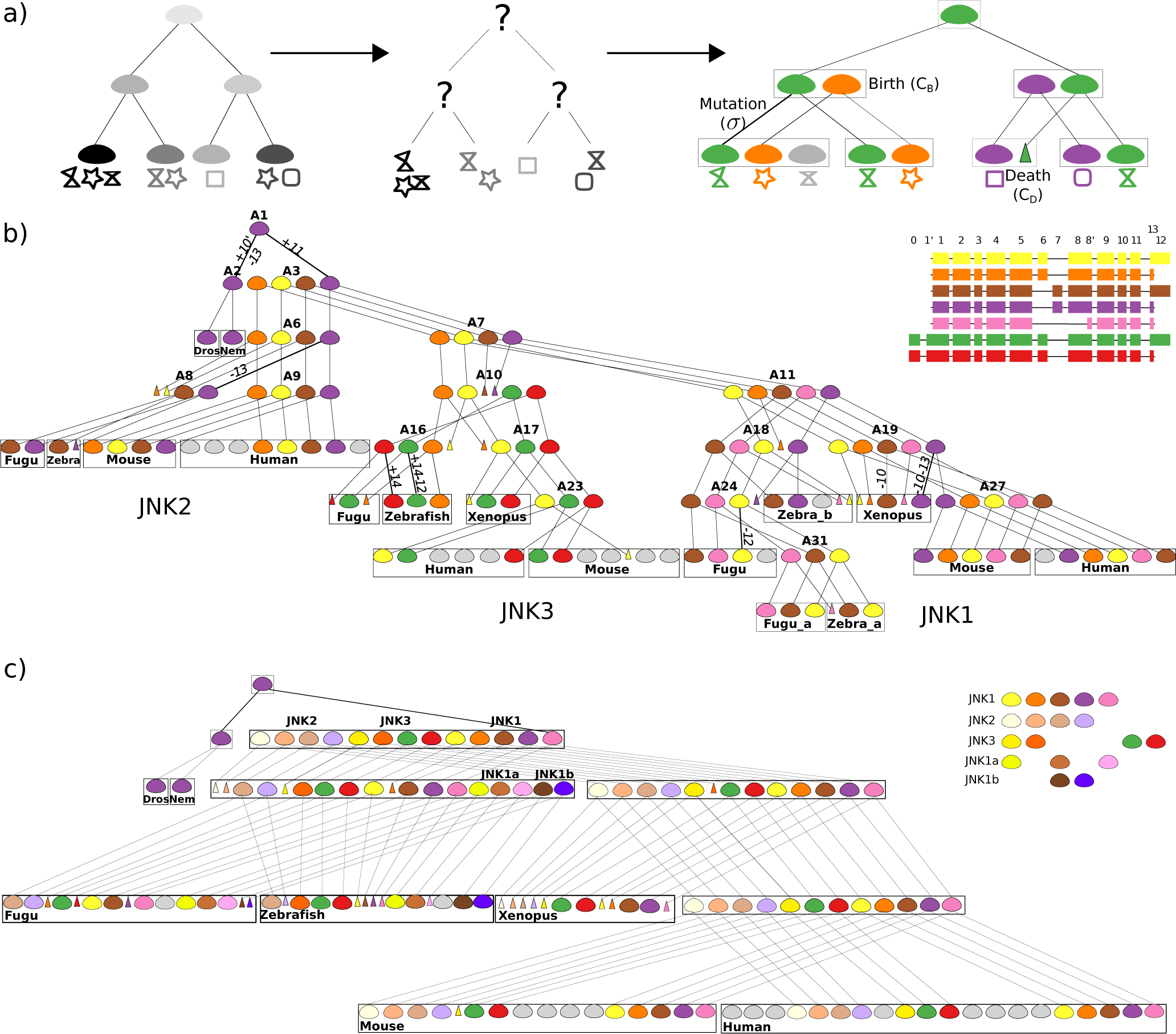
Transcripts’ phylogenies reconstructed by PhyloSofS. **(a)** On the left, example of a phylogenetic gene tree where 8 transcripts (represented by geometrical symbols) are observed in 4 current species (leaves of the tree, colored in different grey tones). These data are given as input to PhyloSofS. In the middle, the problem addressed by PhyloSofS is that of a partial assignment: how to pair transcripts so as to maximize their similarity? On the right, example of a solution determined by PhyloSofS. The transcripts’ phylogeny is a forest comprised of 3 trees (colored differently). The nodes of the input gene tree are subdivided into subnodes corresponding to observed (current) or reconstructed (ancestral) transcripts. The root of a tree stands for the creation of a new transcript and is associated to a cost *C*_*B*_. Triangles indicate transcript deaths and are associated to a cost *C*_*D*_. Mutation events occur along branches and are associated to a cost *σ*. The grey node corresponds to an orphan transcript for which no phylogeny could be reconstructed. **(b)** Transcripts’ phylogeny reconstructed by PhyloSofS for the JNK family. The forest is comprised of 7 trees, 19 deaths (triangles) and 14 orphan transcripts (in grey). Mutation events are indicated on branches by the symbol *+* or *−* followed by the number of the exon being included or excluded (*e.g. +11*). The cost of the phylogeny is 69 (with *C*_*B*_ = 3, *C*_*D*_ = 0 and *σ* = 2). On the top right corner are displayed the exon compositions of the human isoforms for which a phylogeny could be reconstructed. They represent a subset of all the exons composing the 60 transcripts observed in the 7 current species. **(c)** Representation of the transcripts’ phylogeny embedded in the species tree. In this forest, the duplication events are not explicitly indicated, as the different paralogous genes are not linked. There are 2 duplication events giving rise to JNK2 and JNK3 and 2 additional ones for JNK1a and JNK1b (indicated by stars). A given species may contain several paralogous genes. To differentiate the transcripts belonging to the same tree in (b) but to different paralogous genes, different shades of the same colors are used. For example, in human, the transcripts colored in light purple and in purple are issued from JNK2 and JNK1 and belong to the purple tree in (b). For the sake of clarity, mutations are not indicated along the branches. The lists of transcripts appearing in the different genes are displayed in the top right corner.

This case represents a high degree of complexity with 60 observed transcripts assembled from a total of 19 different exons. Most of these transcripts comprise more than 10 exons and the number of transcripts per gene per species varies from 1 to 8 (Fig. 1b-c).

## Results

### Transcripts’ phylogeny for the JNK family

The observed transcripts were collected from the Ensembl [38] database (see *Methods*). PhyloSofS algorithm was run for 10^6^ iterations on the JNK family gene tree and we retained the most parsimonious evolutionary scenario (cost = 69, see *Methods* for a detailed description of the parameters). The reconstructed forest is comprised of 7 transcript trees (Fig. 1b, each tree is colored differently). Each transcript is described as a collection of exons, numbered from *0* to *14* (Fig. 1b, top right corner, and see *Methods* for more details on the numbering). We could reconstruct a phylogeny for 46 out of the 60 observed transcripts. The 14 “orphan” transcripts (leaves in grey) are not conserved across the studied species, and thus likely result in non-functional protein products. Mutations occurring along the branches of the trees are labelled (Fig. 1b, see +/− symbols followed by the number of the included/excluded exon). In total, JNK transcripts’ phylogeny comprises 11 mutations.

The sequences of the JNK genes are highly conserved through evolution (Table 1). While *Drosophila melanogaster* and nematode are the most distant species to human, their unique JNK genes share as much as 78% and 56% sequence identity, respectively, with human JNK1 (Table 1). The sequence identities with human JNK2 and JNK3 are slightly lower (Table 1, in grey). This suggests that the most recent common ancestor of the 7 studied species contained one copy of an ancestral JNK1 gene. Under this assumption, the JNK family gene tree (**S2 Fig, a**) can be reconciled with the species tree (**S2 Fig, b**) by hypothesizing that early duplication events led to the creation of JNK2 and JNK3 in the ancestor common to mammals, amphibians and fishes. JNK1 was then further duplicated in fishes while JNK2 was lost in *Xenopus tropicalis*. A representation of the reconstructed transcripts’ phylogeny embedded in the species tree is displayed on Figure 1c. It permits to appreciate the diversity of transcripts in each species.

**Table 1:**
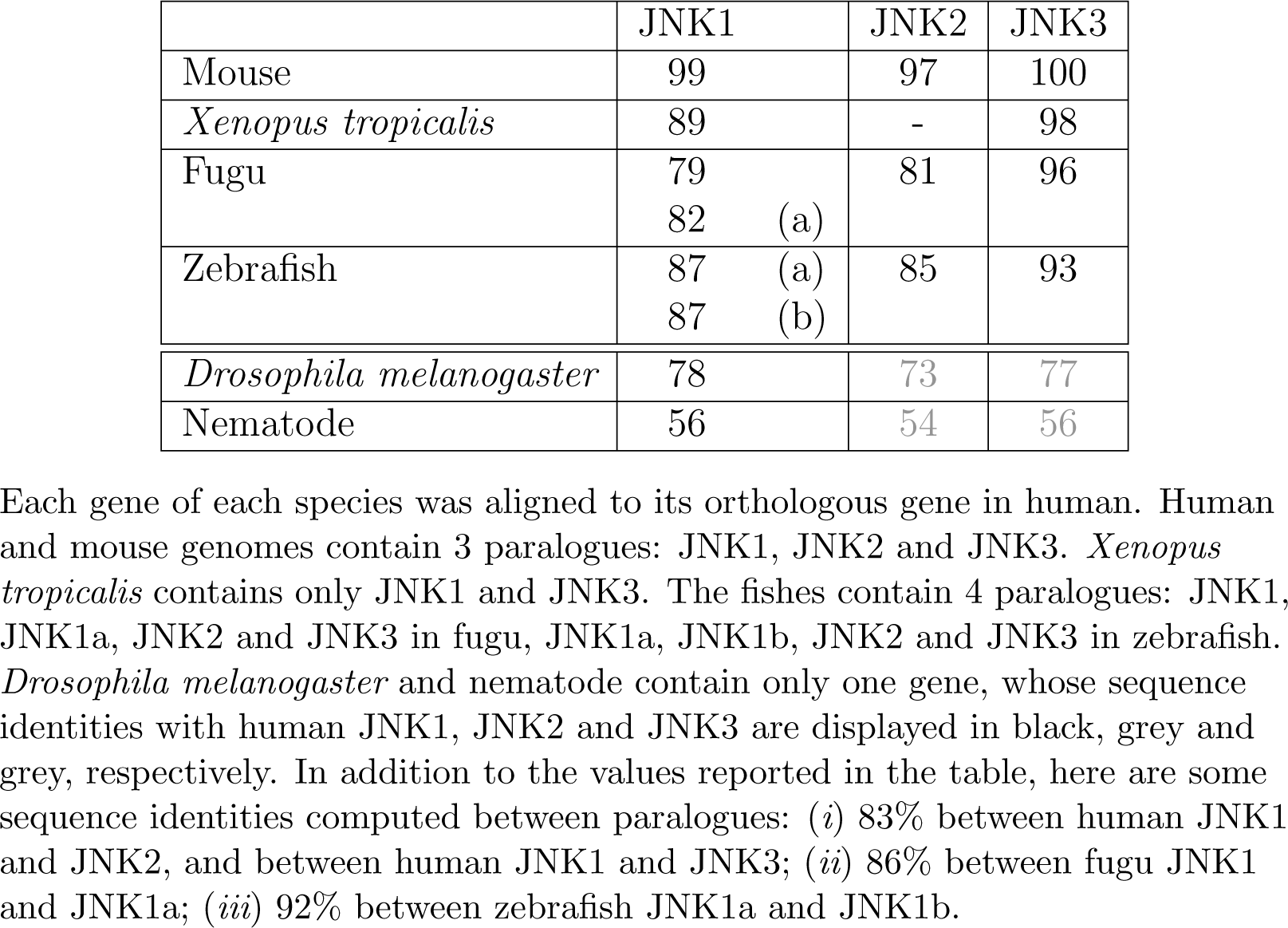
Percentages of sequence identity of JNK genes to human.

|  | JNK1 | JNK2 | JNK3 |
| --- | --- | --- | --- |
| Mouse | 99 | 97 | 100 |
| *Xenopus tropicalis* | 89 | - | 98 |
| Fugu | 79<br>82 (a) | 81 | 96 |
| Zebrafish | 87 (a)<br>87 (b) | 85 | 93 |
| *Drosophila melanogaster* | 78 | 73 | 77 |
| Nematode | 56 | 54 | 56 |

Each gene of each species was aligned to its orthologous gene in human. Human and mouse genomes contain 3 paralogues: JNK1, JNK2 and JNK3. *Xenopus tropicalis* contains only JNK1 and JNK3. The fishes contain 4 paralogues: JNK1, JNK1a, JNK2 and JNK3 in fugu, JNK1a, JNK1b, JNK2 and JNK3 in zebrafish. *Drosophila melanogaster* and nematode contain only one gene, whose sequence identities with human JNK1, JNK2 and JNK3 are displayed in black, grey and grey, respectively. In addition to the values reported in the table, here are some sequence identities computed between paralogues: (i) 83% between human JNK1 and JNK2, and between human JNK1 and JNK3; (ii) 86% between fugu JNK1 and JNK1a; (iii) 92% between zebrafish JNK1a and JNK1b.

The 7 reconstructed trees relate 12 transcripts observed in human across the three genes (Fig. 1b). The transcripts colored the same belong to the same tree and share the same exon composition, even if they come from different gene loci and hence have different amino acid sequences. For instance, the transcript structure including exons *6*, *8* and *12* and excluding exons *0*, *1’*, *7* and *13* (in yellow) is shared by 3 human transcripts present in JNK1, JNK2 and JNK3 (**S2 Fig, c**). Note that this may not be the case in general, for any protein family: the leaves of a tree may have different exon compositions if mutations occur along the branches.

Two pairs of exons, namely *6-7* and *12-13*, are mutually exclusive (**S2 Fig, c**). The associated ASEs can be dated early in the phylogeny (Fig. 1b), before the gene duplication (**S2 Fig, b**). Neither *Drosophila melanogaster* nor nematode contain any of exons *12-13*. Hence, it is equivalent to consider that exon *12* or exon *13* appeared first (compare Fig. 1b and **S9 Fig**). By contrast, exon *7* is clearly predicted as appearing before exon *6* (Fig. 1b, compare purple tree with yellow and orange trees). Noticeably, The two transcripts expressing exons *6* (in orange and yellow) are consistently absent from *Zebrafish* and *Xenopus tropicalis* in JNK1. Although this can correspond to a real loss of transcripts in those species, a more parsimonious explanation would be that the gene annotation in the Ensembl database is incomplete. We searched for direct experimental evidence of the expression of the transcripts in these two organisms using transcriptome sequencing data from hundreds of RNA-Seq libraries (see *Methods*). In *Xenopus tropicalis*, the analysis of exon-exon junctions revealed the expression of transcripts containing a 72bp-long exon with a translated sequence very similar to that of exon *6* in other species. The sequencing support for this exon is strong as it is present in more than two third of the *Xenopus tropicalis* RNA-seq libraries we studied and additionally has evidence from expressed sequence tags (ESTs) data. Additionally, the genomic region corresponding to that exon is strongly conserved (see this link). In *D. rerio*, the analysis of exon-exon junctions also identified one new 72bp-long exon with a translated sequence very similar to exon *6* in other species. Hence, there is significant evidence of the expression of JNK1 transcripts containing exon *6* in both *X. tropicalis* and *D. rerio*, although they are not annotated in Ensembl. This observation gives support to our choice of not penalising death when we reconstruct the transcripts’ phylogeny as a way to account for the incomplete transcript data.

Among the three transcripts appearing after the gene duplication events (Fig. 1b, in pink, green, and red), one transcript features a large deletion encompassing exons *6*, *7* and *8* (JNK1 sub-forest, internal node A11, in pink). Its exon composition is perfectly conserved along the phylogeny (no mutation). We looked for additional RNA-seq support for this transcript and found evidence in 3 Human RNA-seq libraries (out of 166) for reads aligning to the exon-exon junction between exon 7 and 8’. There was no evidence in the RNA-Seq mouse libraries. The two other transcripts are created at the root of the JNK3 sub-forest (ancestor node 10, in green and red). They are characterized by the presence of exons *0* and *1’*, not found in the other paralogues. Another characteristic feature can be observed for the JNK3 gene, namely exon *7* is completely absent from the associated sub-forest. The genomic sequence of exon *7* is present at the JNK3 locus in all species, but it diverged far more in this gene compared to JNK1 and JNK2.

### Mapping of the gene 1D structure onto the protein 3D structure

About eighty structures of human JNKs are available in the PDB (**S1 Table**). This abundance of structural data can be explained by the fact that JNKs are important therapeutic targets and they were crystallized with different inhibitors. The three paralogues share the same fold, which is highly conserved among protein kinases. The structures are highly redundant, with an average root mean square deviation (RMSD) of 1.96 ± 0.71 Å, computed over more than 80% of the protein residues. In order to visualize the correspondence between the gene structure and the protein secondary and tertiary structures, the exons were mapped onto a high-resolution PDB structure (3ELJ [39]) of human JNK1 (Fig. 2, each exon is colored differently). One can observe that the organization of the protein 3D structure is preserved by the 1D structure of the gene. Most of the secondary structures (10 over 12 *α*-helices and 7 over 9 *β*-strands) are completely included in single exons. Moreover, each one of the regions important for the structural stability and/or function of protein kinases (Fig. 2, labelled in black) is included in one single exon (see also **S2 Table**). So are the N-terminal hairpin and the MAPK insert (labelled in grey), two structural motifs specific to the mitogen-activated protein kinase (MAPK) type, to which the JNKs belong. By contrast, binding sites for cofactors and substrates (green circles, see also **S2 Table**) are comprised of residues belonging to different exons. This is expected as binding sites are comprised of segments that can be very far from each other along the protein sequence. Of note, the block formed by exons *1* to *5*, comprising the N-terminal lobe and the A(ctivation)-loop (Fig. 2, from blue to white), is constitutively present in all transcripts belonging to the colored trees on Fig. 1b. The correspondence was also analyzed for the JNK protein from *Drosophila melanogaster* (**S3 Fig, b**). The 3D structures of human JNK1 (**S3 Fig, a**) and *Drosophila melanogaster* JNK (**S3 Fig, b**) are very similar, with a RMSD of 0.68 Å on 251 over 314 (80%) residues. The JNK gene from the *Drosophila melanogaster* genome comprises much fewer exons than the human gene. The match between the borders of these exons and the borders of the secondary structures and known important regions is even better in that species. Considering the high degree of conservation of JNK sequences, one may hypothesize that a good match also exists in all studied species. Our observation is in agreement with a previous study establishing a relationship between exon boundaries and structurally consistent protein regions [19].

**Figure 2.**
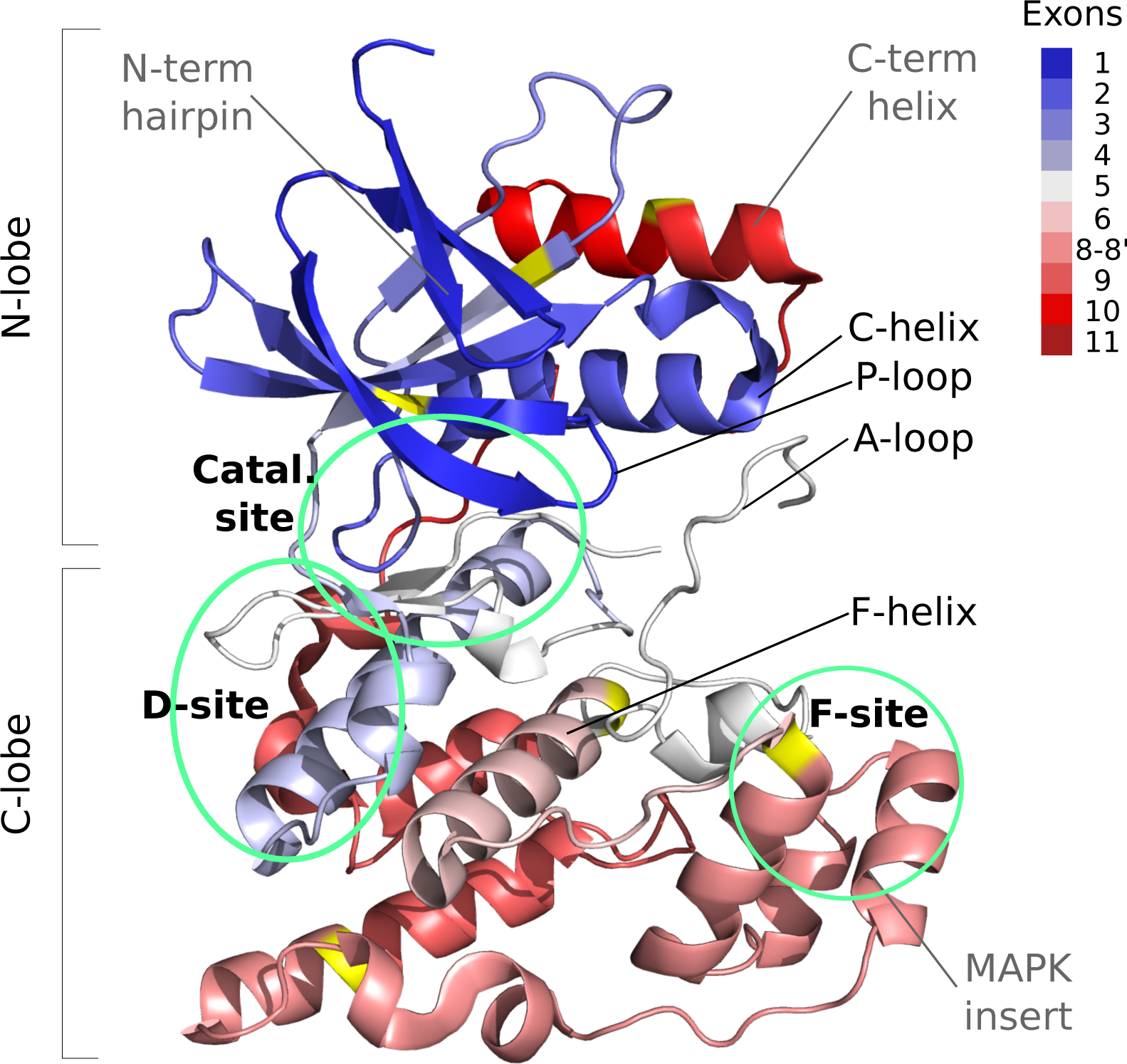
Exons mapped onto the tertiary structure of human JNK1. The protein (residues 7 to 364) is represented as a cartoon and the different exons are colored from blue through white to red. The residues in yellow are at the junction of 2 exons. It should be noted that exons *8* and *8’* used in PhyloSofS actually correspond to only one genomic exon (see *Methods*). The regions labelled in black are common to kinases and were reported in the literature (see [76]) for playing important roles in their structural stability and/or function. The regions labelled in grey are specific to MAP kinases. The green circles indicate the catalytic site and binding sites for JNK cellular partners [63, 44]. The structure was solved by X-ray crystallography at 1.80 Å resolution (PDB code: 3ELJ [39]).

### Properties of the orphan transcripts

We investigated whether the orphan transcripts, for which no phylogeny could be reconstructed (Fig. 1b, grey leaves), displayed peculiar sequence and structural properties compared to the “parented” transcripts (Fig. 1b, colored leaves). Our assumption is that an orphan transcript is less likely to have functional importance. First, the orphan transcripts are significantly smaller than the parented ones (Fig. 3a). While the minimum length for parented transcripts is 308 residues, with an average of 406 ± 40 residues (Fig. 3a, in white), the orphan transcripts can be as small as 124 residues, with an average of 280 ± 88 residues (Fig. 3a, in grey). Second, regarding secondary structure content, both types of transcripts contain about 40% of residues predicted in *α*-helices or *β*-sheets (Fig. 3b). Third, the 3D models generated by PhyloSofS molecular modeling routine for the orphan transcript isoforms are of poorer quality than those for the transcripts belonging to a phylogeny (Fig. 3c-d). The quality of the models was assessed by computing Procheck [40] G-factor and Modeller [41] normalized DOPE score (Fig. 3c-d). A model resembling experimental structures deposited in the PDB should have a G-factor greater than −0.5 (the higher the better) and a normalized DOPE score lower than −1 (the lower the better). The distributions obtained for the parented isoforms are clearly shifted toward better values and are more narrow than those for the orphan transcripts. Finally, the proportion of protein residues being exposed to the solvent (relative accessible surface area *rsa >* 25%) is significantly higher for the orphan isoforms (Fig. 3e), as is the proportion of hydrophobic residues being exposed to the solvent (Fig. 3f). Overall, these observations suggest that simple sequence and structure descriptors enable to distinguish the orphan transcripts from the ones within a phylogeny and that the formers display properties likely reflecting structural instability (large truncations, poorer quality, larger and more hydrophobic surfaces).

**Figure 3.**
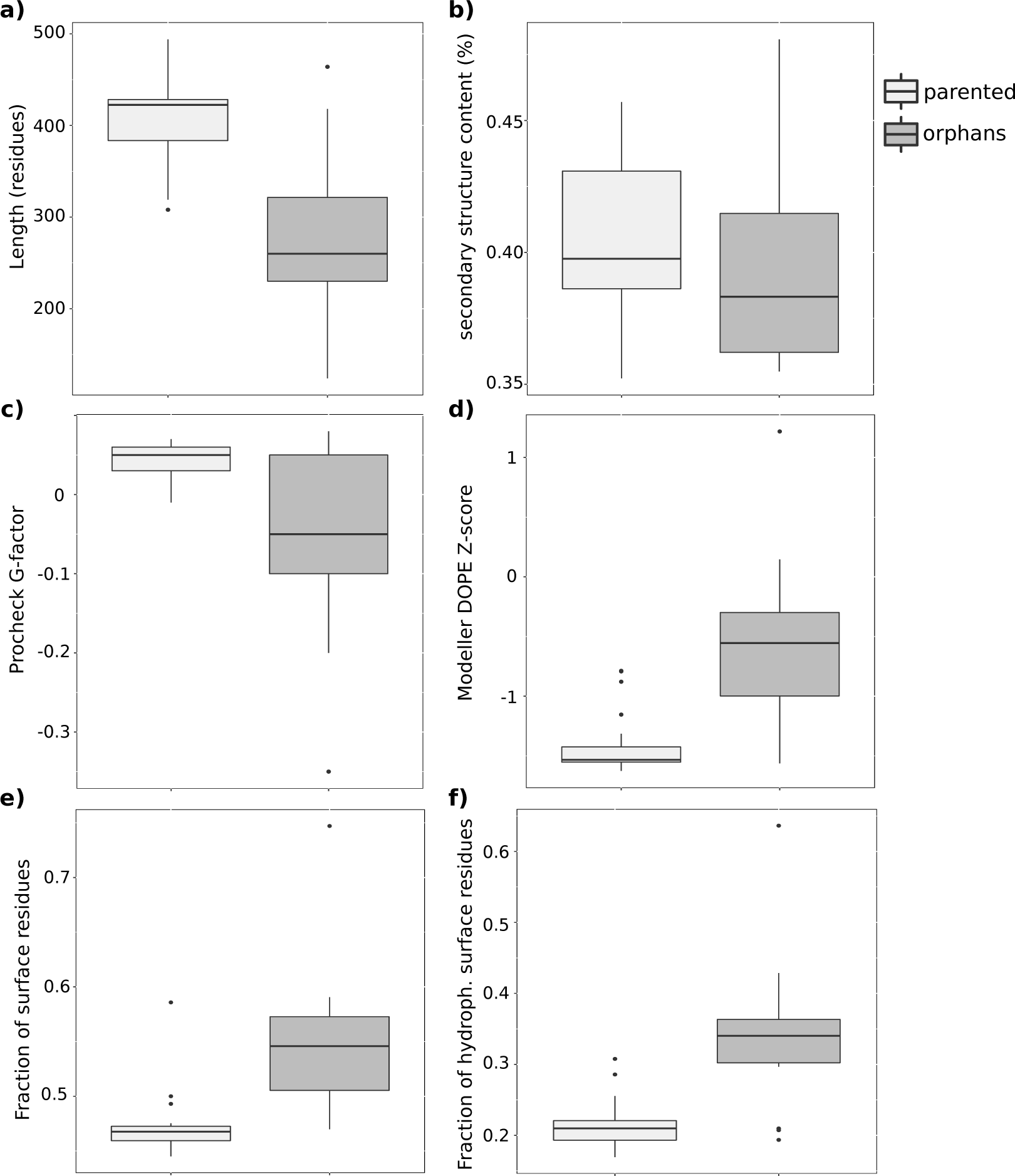
Structural features of the transcript isoforms. Distributions are reported for the parented transcripts (in light gray) and the orphan transcripts (in dark grey) in the transcripts’ phylogeny (see Fig. 1b). **(a)** Length of the transcript (in residues). **(b)** Predicted secondary structure content (in percentages of residues). **(c)** Overall G-factor computed by Procheck [40]. **(d)** Normalized DOPE score computed by Modeller [41]. **(e)** Fraction of protein residues being exposed to the solvent (*rsa >* 0.25). **(f)** Fraction of hydrophobic protein residues being exposed to the solvent (*rsa >* 0.25).

### Subtle changes in the protein’s internal dynamics linked to substrate differential affinity

The two mutually exclusive exons *6* and *7* are particularly important for JNK cellular functions, as they confer substrate specificity. The inclusion or exclusion of one or the other results in different substrate-binding affinities [35, 36]. From a sequence perspective, the two exons are homologous, highly conserved through evolution, and differ only by a few positions (**S4 Fig**). From a structural perspective, they both fold into an *α*-helix, known as the F-helix, followed by a loop (Fig. 2, in light pink). The F-helix was shown to play a central role in the structural stability and catalytic activity of protein kinases [42, 43]. It serves as an anchor for two clusters of hydrophobic residues, namely the catalytic and regulatory spines (see illustration on the PKA kinase on **S5 Fig, a**), and for the HDR motif of the catalytic loop (see illustration on the CDK-substrate complex on **S6 Fig, a**). In the following, we will use these known structural features as proxies for the stability and catalytic competence of the studied isoforms.

The available JNK crystallographic structures and the 3D models generated by PhyloSofS do not display any significant structural change upon exchanging exons *6* and *7*. The catalytic and regulatory spines, together with their anchors in the F-helix, are present in both types of isoforms (**S5 Fig, b-c**). The N-terminal aspartate (D207) of the F-helix, which serves as an anchor for the spines, is 100% conserved in both exons *6* and *7* in the 7 studied species (**S4 Fig**, indicated by an arrow). The two other anchor points are also present, namely I214 and L/M218 (**S4 Fig**, indicated by arrows). Moreover, the characteristic H-bond pattern with the HRD motif and the associated strained backbone conformation are also observed in both types of isoforms (**S6 Fig, b-c**). Consequently, both exons *6* and *7*, and thus the isoforms containing them, possess the structural features known to be important for kinase catalytic activity and/or regulation.

To further investigate the potential impact of the inclusion/exclusion of exon *6* or *7* on the dynamical behavior of the protein, we performed all-atom molecular dynamics (MD) simulations of the human isoforms colored in orange and purple on Figure 1b. We shall refer to these isoforms as JNK1*α* (with exon *6*) and JNK1*β* (with exon *7*), in agreement with the nomenclature found in the literature [35]. JNK1*α* and JNK1*β* were simulated in explicit solvent for 250 ns (5 replicates of 50 ns, see *Methods*). The backbone atomic fluctuation profiles of the two isoforms are very similar (Fig. 4a, orange and purple curves), except for the A-loop which is significantly more flexible in JNK1*α*: the region from residue 176 to 188 displays averaged C*α* fluctuations of 1.55 ± 0.28 Å in JNK1*α* and of 0.98 ± 0.16 Å in JNK1*β* (Fig. 4a). We should stress that this loop displays the highest deviations among the JNK structures available in the PDB and often comprises unresolved residues. The two exons, *6* and *7*, have similar backbone flexibility. In the F-helix, the anchor residues for the spines, D207, I214 and M218 adopt stable and very similar conformations (Fig. 4b). Moreover, the HRD backbone strain and the associated H-bond pattern are maintained along the simulations of both systems (**S8 Fig, a-b**). Consequently, the observations realized on the static 3D models hold true when simulating their dynamical behavior: the *6* /*7* variation does not induce any drastic change on the protein’s overall shape and behaviour.

**Figure 4.**
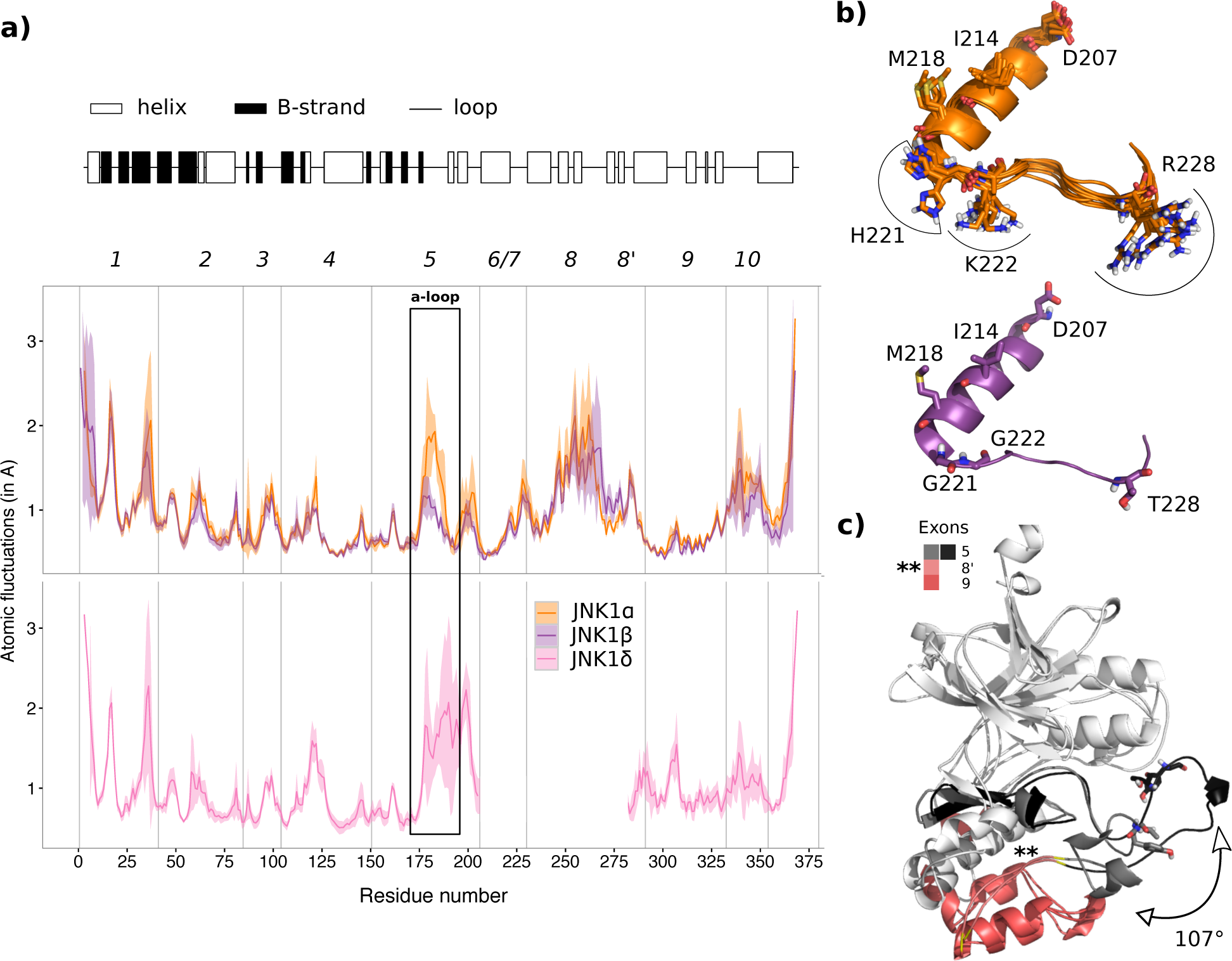
Dynamical behavior of the human JNK1 isoforms in solution. **(a)** The secondary structures for JNK1*α* (with exon *6*) are depicted on top (the profiles for the 2 other isoforms are very similar, see **S7 Fig**). The atomic fluctuations (computed on the C*α*) averaged over 5 50-ns MD replicates are reported for JNK1*α* in orange, JNK1*β* in purple and JNK1*δ* in pink. The envelopes around the curves indicate the standard deviation. **(b)** Representative MD conformations obtained by clustering based on position 228 (RMSD cutoff of 1.5 Å). There are 8 conformations for JNK1*α* (in orange) and only 1 for JNK1*β* (in purple). **(c)** Superimposed pair of MD conformations illustrating the amplitude of the A-loop motion in JNK1*δ* (see *Materials ad Methods* for details on the calculation of the angle). Exons *5*, *8’* and *9* are indicated by colors and labels. For clarity, *8’* is also indicated by two stars on the structure.

Nevertheless, we observe differences in the side-chain flexibilities of a few residues lying in the loop following the F-helix between the two isoforms (Fig. 4b). On the one hand, in exon *6* (in orange), the polar and positively charged residues H221, K222 and R228 are exposed to the solvent and display large amplitude side-chain motions. These amino acids are 100% conserved in exon *6* across all species (**S4 Fig**). On the other hand, in exon *7* (Fig. 4b, in purple), G221, G222 and T228 have small side chains with much reduced motions. While G221 is conserved across all species, position 222 is variable and position 228 features G, T or S (**S4 Fig**). This region of the protein is involved in the binding of substrates (see Fig. 2, F-site). Moreover, in both isoforms, we predicted residues 223-230 as directly interacting with cellular partners (see *Methods*). Consequently, one may hypothesize that the differences highlighted here may be crucial for substrate molecular recognition specificity. The positive charges, high fluctuations, high solvent accessibility and high conservation of residues H221, K222 and R228 in JNK1*α* support a determinant role for these residues in selectively recognizing specific substrates.

### Structural dynamics of a newly identified isoform

Our reconstruction of the JNK transcripts’ phylogeny highlighted a JNK1 isoform (Figure 1b, in pink) that has not been documented in the literature so far. It is expressed in human, mouse and fugu fish (Figure 1b), suggesting that it could play a functional role in the cell. To investigate this hypothesis, we analyzed the 3D structure and dynamical behavior of this isoform in human. We refer to it as JNK1*δ*.

JNK1*δ* displays a large deletion (of about 80 residues), lacking exons *6*, *7* and *8*. It does not contain the F-helix, shown to be crucial for kinases structural stability [42], nor the MAPK insert, involved in the binding of the phosphatase MKP7 [44] (Fig. 2). The 3D model generated by PhyloSofS superimposes well to those of JNK1*α* and JNK1*β*, with a RMSD lower than 0.5 Å on 245 residues. This is somewhat expected as we use homology modeling. Nevertheless, cases were reported in the literature where homology modeling detected big changes in protein structures induced by exon skipping [45]. In the model of JNK1*δ*, the F-helix present in JNK1*α* and JNK1*β* (residues 207 to 220) is replaced by a loop (residues 282 to 288) corresponding to exon *8’* (Fig. 4c, indicated by the two stars). The sequence of this loop (exon *8’*) does not share any significant identity with the F-helix (N-terminal parts of exons *6* and *7*), except for the N-terminal residue which is an aspartate, namely D282 (D207 in JNK1*α* and JNK1*β*). This replacement results in the regulatory spine being intact in JNK1*δ* (**S5 Fig, d**, in red). Moreover, the HRD motif’s strained backbone conformation and the associated H-bond pattern, which are stabilized by the aspartate, are maintained (**S6 Fig, d**). By contrast, the catalytic spine lacks its two anchors (**S5 Fig, d**, in yellow).

JNK1*δ* was simulated in explicit solvent for 250 ns (5 replicates of 50 ns). The isoform displays stable secondary structures (**S7 Fig**, at the bottom) and atomic fluctuations comparable to those of JNK1*α* and JNK1*β* (Fig. 4a, pink curve to be compared with the purple and orange curves). The C*α* atomic fluctuations averaged over the loop replacing the F-helix are of 0.88 ± 0.18 Å. This is higher than the values computed for the F-helix in JNK1*α* and JNK1*β* (0.57 ± 0.10 Å and 0.53 ± 0.09 Å), but it still indicates a limited flexibility. Moreover, the N-terminal aspartate D282 establishes stable H-bonds with the HRD motif along all but one of the replicates (**S8 Fig, a**, on the right) and the HRD motif’s backbone remains in a strained conformation (**S8 Fig, b**, on the right), as was observed for JNK1*α* and JNK1*β*. Consequently, JNK1*δ* seems stable in solution, and, as observed on the static 3D model, the absence of the F-helix in this isoform is partially compensated by the presence of D282, which is sufficient to maintain H-bonds with the HRD motif and a resulting backbone strain of the motif, important for kinase structural stability.

The main difference between JNK1*δ* and the two other isoforms lies in the amplitude of the motions of the A-loop. In JNK1*δ*, the C-terminal part of the A-loop can detach from the rest of the protein along the simulations (Fig. 4c). The amplitude of the angle computed between the most retracted conformation (in grey) and the most extended one (in black) is 107°. By contrast, in JNK1*α* and JNK1*β*, the A-loop always stays close to the rest of the protein, with amplitude angles of 18° and 19°, respectively. The A-loop contains two residues, T183 and Y185 (Fig. 4c, highlighted in sticks), whose phosphorylation is required for JNK activation. We hypothesize that the large amplitude motion in JNK1*δ* might favor their accessibility and, in turn, the activation of the protein.

### Set of equivalent transcripts’ phylogenies

The size of the search space for the transcripts’ phylogeny reconstruction grows exponentially with the number of observed transcripts (leaves). To explore that space, the heuristic algorithm implemented in PhyloSofS relies on a multi-start iterative procedure and on the computation of a lower bound to early filter out unlikely scenarios (see *Methods*). Depending on the input data and the set of parameters, it may find several solutions with equivalent costs. Over 10^6^ iterations of the program, the forest described above (Fig. 1b, or **S9 Fig** with branch swapping), comprising 7 trees, 19 deaths and 14 orphans, was visited 1 219 times. An alternative phylogeny was visited 310 times, that comprises the same number of trees and orphans, but 2 more deaths (**S10 Fig**). The difference between the two forests lies among the fugu JNK1 transcripts, where one transcript belongs to the orange tree (**S10 Fig**) instead of the yellow one (Fig. 1b). The two trees differ by the inclusion or exclusion of exon *12* or *13*, and the re-assigned transcript lacks both exons. Consequently, the new branching results in the loss of exon *13* between the internal nodes A11 and A18 (**S10 Fig**), instead of the loss of exon *12* between A24 and fugu JNK1 (Fig. 1b). Another forest with the same cost comprising 8 trees, 23 deaths and 13 orphans was visited 190 times (**S11 Fig**). The additional tree is created in the internal node A10 and links two observed JNK3 transcripts: one from the mouse that was previously orphan (Fig. 1b) and one from zebrafish that previously belonged to the green tree. Both transcripts are truncated at the C-terminus and lack exons *12* and *13*. Consequently, this new branching avoids the loss of exon *12* between A16 and zebrafish JNK3. Overall the differences between the three solutions are minor and these ambiguities do not impact our interpretation of the results.

### Unresolved residues in the 3D models

In the 3D models generated by PhyloSofS, the N-terminal exons *0* and *1’* and the C-terminal exons *12* and *13* are systematically missing. This is due to the lack of structural templates for these regions. Using a threading approach instead of PhyloSofS homology modeling routine (see *Methods*) did not enable to improve their reconstruction. In fact, the models generated by the threading algorithm are very similar to those generated by PhyloSofS.

At the C-terminus, exons *12* and *13* are completely predicted as intrinsically disordered (**S12 Fig, a** and **S12 Fig, b**, blue curve). At the N-terminus, exons *0* and *1’* contain two segments of about 10 residues predicted as disordered protein-binding regions (**S12 Fig, b**, orange curve), *i.e* regions unable to form enough favorable intra-chain interactions to fold on their own and likely stabilized upon interaction with a globular protein partner [46]. These exons are present in only two JNK3 transcript isoforms (Fig. 1b, colored in red and green).

## Discussion

To what extent the transcript diversity generated by AS translates at the protein level and has functional implications in the cell remains a very challenging question and has been subject to much debate [47, 48]. The present work contributes to elaborating strategies to answer it, by crossing sequence analysis and phylogenetic inference with molecular modeling. We report the first joint analysis of the evolution of alternative splicing across several species and of its structural impact on the produced isoforms. The analysis was performed on the JNK family, which represents a high interest for medicinal research and for which a number of human isoforms have been described and biochemically characterized.

Firstly, our results allowed dating an ASE consisting of two mutually exclusive homologous exons (*6* and *7*) in the ancestor common to mammals, amphibians and fishes. We find that the most ancient of these two exons is exon *7*. By characterizing in details the structural dynamics of two human isoforms, JNK1*α* and JNK1*β*, bearing one or the other exon, we could emphasize subtle changes associated to this ASE and identify residues that may be responsible for the selectivity of the JNK isoforms toward their substrates. Secondly, we highlighted an isoform that was not previously described in the literature, namely JNK1*δ*. Despite displaying a large deletion (about 80 residues), it is conserved across several species and short MD simulations suggest that it is stable in solution. According to the APPRIS database v20 [24], there are 4 peptides matching this isoform in publicly available proteomics data. By comparison, the other human JNK1 isoforms with a phylogeny have between 5 and 7 matching peptides, while the orphan transcripts identified by our analysis have between zero and 2 matching peptides, suggesting that JNK1*δ* is indeed translated and stable in solution. Hence, considering that the catalytic site is intact in JNK1*δ*, we propose that this isoform might be catalytically competent and that the large amplitude motion of the A-loop observed in the simulations might facilitate the activation of the protein by exposing a couple of tyrosine and threonine residues that are targeted by MAPK kinases. The validation of this hypothesis would require further calculations and experiments that fall beyond the scope of this study. Already, this interesting result suggests that our approach could be used to identify and characterize new isoforms, that may play a role in the cell and thus serve as therapeutic targets. Thirdly, we found characteristics specific to the JNK3 isoforms, namely the absence of exon *7* and the presence of twos exons (*0* and *1’*) containing regions predicted to be disordered and involved in interactions. These observations suggest specific competences or functions for this gene. Studies investigating the gain/loss of alternative splice forms associated to gene duplication at large scale [49, 50] have highlighted a wide diversity of cases and have suggested that it depends on the specific cellular context of each gene. Although we did not have a sufficient sample resolution to confirm it with RNA-Seq data, JNK3 is reported to be specifically expressed in the heart brain and testes [35]

Our approach enables to go beyond a description of transcript variability across species and/or across genes. Indeed, by reconstructing phylogenies, we do not only cluster transcripts but we also add a temporal dimension to the analysis. To efficiently search the space of possible phylogenies, the algorithm implemented in PhyloSofS relies on a multi-start iterative procedure and on the computation of a lower bound that enables to early eliminate unsuitable candidate solutions (see *Methods*). For the JNK family, the execution of 1 million iterations took about two weeks on a single CPU. This case represents a high level of complexity as most of the transcripts contain more than 10 exons (the average number of exons per gene being estimated at 8.8 in the human genome [51]) and up to 8 transcripts are observed within each species (it is estimated that about 4 distinct-coding transcripts per gene are expressed in human [24]). To reduce the computing time, the user can easily parallelize the multi-start iterative search on multiple cores and s/he has the possibility to give as input a previously computed value for the lower bound (to increase the efficiency of the cut). This implementation makes the reconstruction of transcripts’ phylogenies feasible for any gene family. We should stress that the problem of pairing transcripts across homologous and paralogous genes between different species, addressed here, is much more complex than that of inferring the transcripts’ phylogeny of each gene separately. This is because reconciliation between gene and species tree is needed, the homologous exons are evolutionary more distant and the problem size increases.

Our phylogenies may be impacted by two main sources of error coming from the input data. Specifically, under-annotation of transcripts can lead to missing distant evolutionary relationships. To deal with this issue, we set the cost associated to transcript death to zero. This enables to construct trees that can relate transcripts possibly very far from each other in the phylogeny (*i.e.* expressed in very distant species, because some species in between are under-annotated). This parameter may be tuned by the user depending on the quality and reliability of the input data. A second source of error comes from annotated transcripts that are not translated or not functional at the protein level. However, we do not expect that these transcripts will significantly pollute the phylogenetic reconstruction. Indeed, they are likely not conserved across species and thus will be attributed the status of orphans in the phylogenetic reconstruction. Moreover, we have emphasized an independent source of evidence coming from their structural characterization which can help us flag them. The reliability of the transcript expression data clearly constitutes a present limitation of the method. However, as experimental evidence accumulate and precise quantitative data become available, computational methods such as PhyloSofS will become instrumental in assessing the contribution of AS in protein evolution.

Although PhyloSofS was applied here to study the evolution of transcripts in different species, it has broad applicability and can be used to study transcript diversity and conservation among diverse biological entities. The entities could be at the scale of (*i*) one individual/species (tissue/cell differentiation), (*ii*) different species (matching cell types), (*iii*) population of individuals affected or not by a multifactorial disorder. In the first case, the tree given as input should describe checkpoints during cell differentiation and PhyloSofS will provide insights on the ASEs occurring along this process. In the second case, PhyloSofS can be applied to study one particular tissue across several species in a straightforward manner (explicitly dealing with the dimension of different tissues requires further development). In the third case, the tree given as input may be constructed based on genome comparison, a biological trait or disease symptoms. PhyloSofS can be used to evaluate the pertinence of such criteria to relate the patients, with regards to the likelihood (parsimony) of the associated transcripts scenarios. This case is particularly relevant in the context of medical research.

## 1. Methods

### 1.1. PhyloSofS workflow

PhyloSofS can be applied to single genes or to gene families. The input is a binary tree (called a gene tree) describing the phylogeny of the gene(s) of interest for a set of species (Fig. 1a, on the left), and the ensemble of transcripts observed in these species (symbols at the leaves). PhyloSofS comprises two main steps:

a. It reconstructs a forest of phylogenetic trees describing plausible evolutionary scenarios that can explain the observed transcripts (Fig. 1a, on the right). The forest is embedded in the input gene tree and is reconstructed by using the maximum parsimony principle. The root of a tree corresponds to the creation of a new transcript, each leaf stands for an observed transcript and a dead end (indicated by a triangle on Fig. 1a, on the right) indicates a transcript loss. Transcripts can mutate along the branches of the trees.
b. It predicts the three-dimensional structures of the protein isoforms corresponding to the observed transcripts by using homology modeling. The 3D models are then annotated with quality measures and with exon labels.

PhyloSofS comes with helper functions for the visualization of the output transcripts’ phylogeny(ies) and of the isoforms’ molecular models. The program is implemented in Python 3 and freely available at GitHub under MIT license: https://github.com/PhyloSofS-Team/PhyloSofS.

#### Step a. Transcripts’ phylogenies reconstruction

For simplicity and without loss of generality, we describe here the case of one gene of interest studied across several species. The gene is represented by an ensemble *E* of *n*_*e*_ exons. The identification and alignment of the *n*_*e*_ homologous exons between the different transcripts must be performed prior to the application of the method (see below for details on data preprocessing for the JNK family). The *n*_*s*_ transcripts of species *s* are described by a binary table *T*^*s*^ of *n*_*e*_ × *n*_*s*_ elements, where 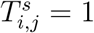 if exon *i* is included in transcript *j* (**Fig. S1 Fig a**, see colored squares), 0 if it is excluded (white squares).

We model transcripts evolution as a two-level process, at the gene and transcript levels, as described by Christinat and Moret [15]. At the level of the gene, each exon can be either absent, alternative or constitutive. This status is inferred from the occurrence of the exon in the transcripts. Hence, for a given species *s*, a vector *g*^*s*^ of length *n*_*e*_ encodes the state of each exon by the values {0, 1, 2} for absent, alternative and constitutive, respectively (**Fig. S1 Fig b**, white, black/white and black squares). At the leaves (current species), the components of *g*^*s*^ are calculated as:

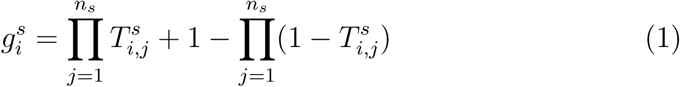

As in [15], the *g*^*s*^ vectors for internal nodes (ancestral species) are determined by using Sankoff’s algorithm [52]. Dollo’s parsimony principle is also respected, such that an exon cannot be created twice [53]. If different exon states have equal cost, we follow the priority rule 2 *>* 0 *>* 1.

Three evolutionary events are considered, namely creation, death and mutation of a transcript with costs *C*_*B*_, *C*_*D*_ and *σ*, respectively. The mutation cost *σ* is accounted for only when the associated evolutionary change occur at the level of the transcript (Table 2). This reflects the fact that changes at the level of the gene affects the expression of exons in the transcripts but changes at the level of the transcripts do not affect the gene structure. For instance, if an exon is absent in a parent and becomes present in the child, then this change of status at the transcript level will be penalized by *σ* only if the exon could be absent in the child, *i.e.* its status at the gene level is “alternative”. If the “constitutive” exon is the child, then the mutation is not penalized (Table 2, compare the cells (0,0)*→*(1,1) and (0,0)*→*(1,2).

**Table 2:**
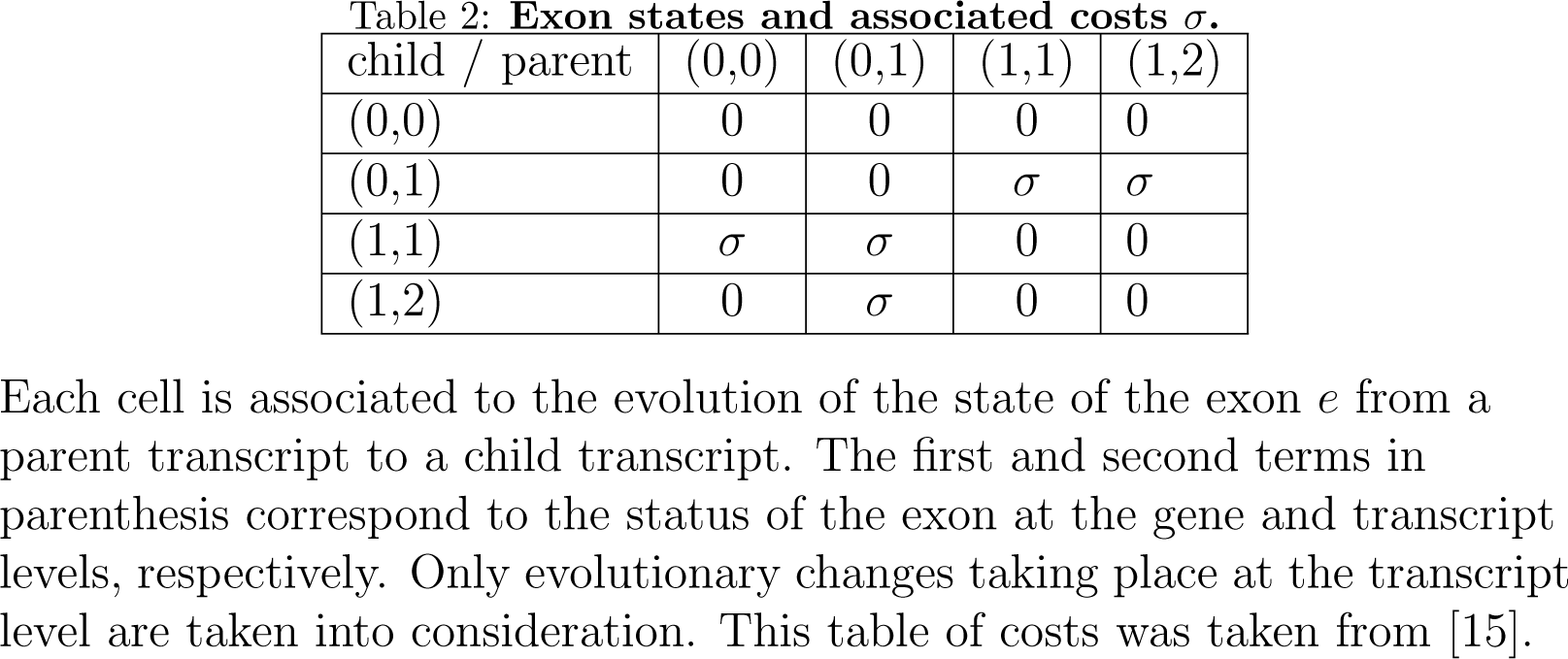
Exon states and associated costs *σ*.

Each internal node of the gene tree, representing an ancestral species, is expanded in several subnodes, representing the transcripts of the gene in this ancestral species (**Fig. S1 Fig c**). There exist three types of subnodes: binary (two transcript children), left (one transcript child in the node’s left child) and right (one transcript child in the node’s right child). Left and right subnodes imply that a transcript death occurred along the branch. A *forest structure S* is fixed by setting *n*_*b*_, *n*_*l*_ and *n*_*r*_ the respective numbers of binary, left and right subnodes for every internal node of the gene tree. The cost associated to structure *S* is calculated as *C*_*S*_ = *C*_*birth*_(*S*) + *C*_*death*_(*S*), where *C*_*birth*_(*S*) and *C*_*death*_(*S*) are the total costs of creation and loss of transcripts, expressed as:

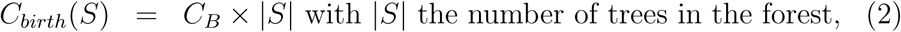

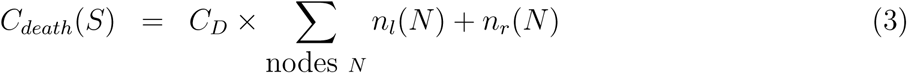

Given a forest structure, a *transcripts’ phylogeny* determines the pairings of transcripts at each internal node (**Fig. S1 Fig d**). The cost of the transcripts’ phylogeny *φ* complying with the forest structure *S* is calculated as:

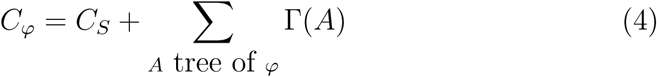

where Γ(*A*) is computed for each tree *A* of *φ* by evaluating the changes of exon states along the branches of *φ*:

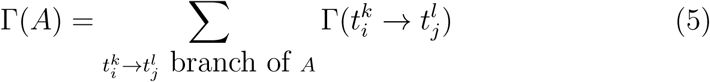

where 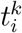 is the parent transcript, *i*^*th*^ subnode of node *k*, 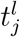 is the child transcript, *j*^*th*^ subnode of node *l* and 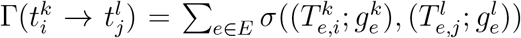, with 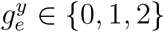 the state of exon *e* at the level of the gene at node *y* and 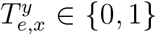 the state of exon *e* at the level of the *x*^*th*^ transcript of node *y*. The evolution costs *σ* are given in Table 2.

PhyloSofS algorithm seeks to determine the scenario with the smallest number of evolutionary events, *i.e.* the transcripts’ phylogeny with the minimum cost (**Fig. S1 Fig c-d**). It proceeds as follows:

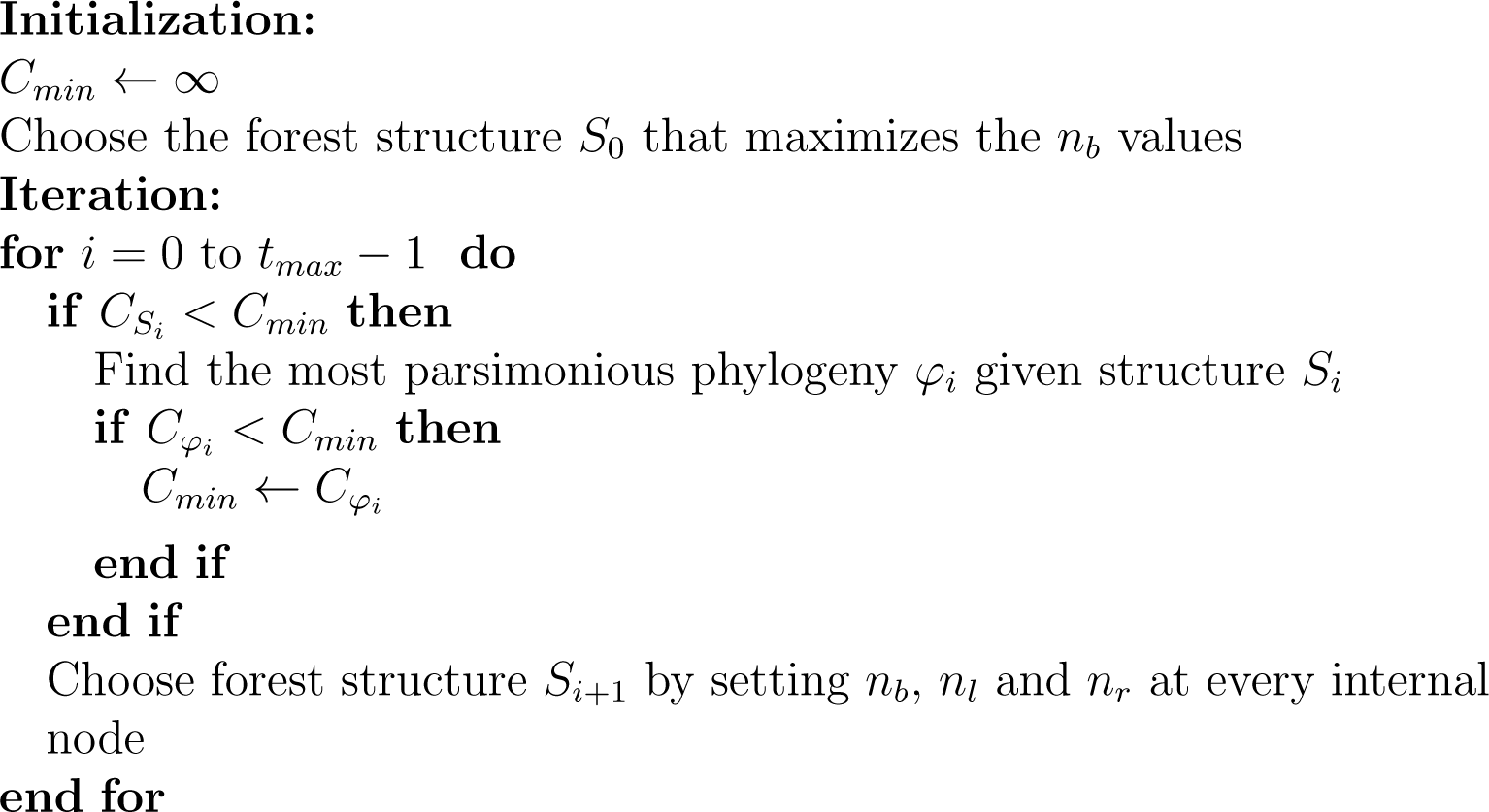

To efficiently search the space of all possible forest structures (**Fig. S1 Fig c**), PhyloSofS relies on a multi-start iterative procedure. Random jumps in the search space are performed until a suitable forest structure *S*_*i*_ (with 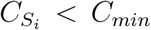) is found. The cost 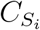 of the forest structure *S*_*i*_ serves as a lower bound for the cost 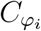 of the phylogeny *φ*_*i*_. Forest structures that are too costly are simply discarded, without calculating the corresponding phylogenies. As the algorithm finds better and better solutions, the cut becomes more and more efficient. The phylogeny *φ*_*i*_ is reconstructed by using dynamic programming. Sankoff’s algorithm is applied bottom up to compute the minimum pairing costs between transcripts (**Fig. S1 Fig d**, each transcript is represented by a matrix of costs). At each internal node, the pairings are determined by using a specific version of the branch-and-bound algorithm [54] (see *Supplementary Text S1*). If the reconstructed phylogeny is more parsimonious than those previously visited (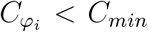), then the minimum cost *C*_*min*_ is updated. There may be more than one phylogeny with minimum cost that comply with a given structure *S*_*i*_. The next forest structure *S*_*j*_ will be randomly chosen among the immediate neighbors of *S*_*i*_ (**Fig. S1 Fig d**). Two structures are immediate neighbors if each one of them can be obtained by an elementary operation applied to only one node of the other one (**S13 Fig**). If the phylogeny *φ*_*j*_ is such that 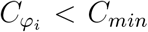, then the next forest structure will be chosen among the neighbors of *S*_*j*_, which serves as a new “base” for the search. Otherwise, the algorithm continues to sample the neighborhood of *S*_*i*_. This step-by-step search is applied until no better solution can be found. At this point, a new random jump is performed. The total number of iterations *t*_*max*_ is given as input by the user (1 by default).

We should stress that PhyloSofS algorithm is designed to deal with much more complex cases than those reported in [15] in a computationally tractable way. Hence, it differs from the algorithm reported in [15] in several respects. First, our multi-start iterative strategy relies on random jumps in the forest structure space combined with systematic local exploration around the best current solution, while Christinat and Moret [15] proposed an exhaustive generation and evaluation of forest structures. Secondly, we have designed a branch-and-bound algorithm specifically adapted to the problem of determining the best phylogeny complying with a given forest structure (see *Supplementary Text S1*). Both aspects contribute to PhyloSofS efficiency in reconstructing transcripts’ phylogenies.

PhyloSofS generates PDF files displaying the computed transcripts’ phylogenies using a Python driver to the Graphviz [55] DOT format.

#### Step b. Isoforms structures prediction

The molecular modeling routine implemented in PhyloSofS relies on homology modeling. It takes as input an ensemble of multi-fasta files (one per species) containing the sequences of the splicing isoforms. For each isoform, it proceeds as follows:

1. search for homologous sequences whose 3D structures are available in the PDB (templates) and align them to the query sequence;
2. select the *n* (5 by default, adjustable by the user) best templates;
3. build the 3D model of the query;
4. remove the N- and C-terminal residues unresolved in the model (no structural template);
5. annotate the model with sequence and structure information.

Step 1 makes extensive use of the HH-suite [37] and can be decomposed in: (a) search for homologous sequences and building of a multiple sequence alignment (MSA), by using HHblits [56], (b) addition of secondary structure predictions, obtained by PSIPRED [57], to the MSA, (c) generation of a profile hidden markov model (HMM) from the MSA, (d) search of a database of profile HMMs for homologous proteins, using HHsearch [58]. Step 3 is performed by Modeller [41] with default options. Step 5 consists in: (a) inserting the numbers of the exons in the *β*-factor column of the PDB file of the 3D model, (b) computing the proportion of residues predicted in well-defined secondary structures by PSIPRED [57], (c) assessing the quality of the model with Procheck [40] and with the normalized DOPE score from Modeller, (d) determining the by-residue solvent accessible surface areas with Naccess [59] and computing the proportions of surface residues and of hydrophobic surface residues.

### 1.2. Retrieval and pre-processing of JNK annotated transcriptome data

The peptide sequences of all splice variants from the JNK family observed in human, mouse, *Xenopus tropicalis*, zebrafish, fugu, *Drosophila melanogaster* and nematode were retrieved from Ensembl [38] release 84 (March 2016) along with the phylogenetic gene tree. Only the transcripts containing an open reading frame and not annotated as undergoing nonsense mediated decay or lacking 3’ or 5’ truncation were retained.The isoforms sharing the same amino acid sequence were merged. The homologous exons between the different genes in the different species were identified by aligning the sequences with MAFFT [60], and projecting the alignment on the human annotation. They do not necessarily represent exons definition based on the genomic sequence and this can be explained by two reasons. First, the gene structure may be different from one species to another. For instance, the third and fourth exons of human JNK1 genes are completely covered by a single exon in the *Drosophila melanogaster* JNK gene (**S3 Fig**). In cases like this, we keep the highest level of resolution and define two exons (*e.g.* numbered *3* and *4*). Secondly, it may happen that a transcript contains only a part of an exon in a given species translated in another frame. In that case, we define two exons sharing the same number but distinguished by the prime symbol (*e.g.* exons *8* and *8’*). In total, 64 transcripts comprised of 38 exons were given as input to PhyloSofS.

### 1.3. PhyloSofS’ parameter setting

To set the parameters, two criteria were taken into consideration. First, the different genomes available in Ensembl are not annotated with the same accuracy and the transcriptome data and annotations may be incomplete. This may challenge the reconstruction of transcripts’ phylogenies across species. To cope with this issue, we chose not to penalize transcript death (*C*_*D*_=0). Second, the JNK genes are highly conserved across the seven studied species (Table 1), indicating that this family has not diverged much through evolution. Consequently, we set the transcript mutation and birth costs to *σ* = 2 and *C*_*B*_ = 3 (*C*_*B*_ < *σ* × 2). This implies that few mutations will be tolerated along a phylogeny. Prior to the phylogenetic reconstruction, PhyloSofS removed 19 exons that appeared in only one transcript (default option), reducing the number of transcripts to 60. This pruning enables to limit the noise contained in the input data and to more efficiently reconstruct phylogenies. PhyloSofS algorithm was then run for 10^6^ iterations.

The 3D models of all observed isoforms were generated by PhyloSofS molecular modeling routine by setting the number of retained best templates to 5 (default parameter) for every isoform.

### 1.4. Analysis of JNK tertiary structures

The list of experimental structures deposited in the PDB for the human JNKs was retrieved from UniProt [61]. The structures were aligned with PyMOL [62] and the RMSD between each pair was computed. Residues comprising the catalytic site were defined from the complex between human JNK3 and adenosine mono-phosphate (PDB code: 4KKE, resolution: 2.2 Å), as those located less than 6 Å away from the ligand. Residues comprising the D-site and the F-site were defined from the complexes between human JNK1 and the scaffolding protein JIP-1 (PDB code: 1UKH, resolution: 2.35 Å [63]) and the catalytic domain of MKP7 (PDB code: 4YR8, resolution: 2.4 Å [44]), respectively. They were detected as displaying a change in relative solvent accessibility *>*1 Å^2^ upon binding.

The I-TASSER webserver [64, 65, 66] was used to try and model the regions for which no structural templates could be found. DISOPRED [67] and IUPred [68] were used to predict intrinsic disorder. JET2 [69] was used to predict binding sites at the surface of the isoforms.

### 1.5. Molecular dynamics simulations of human isoforms

The 3D coordinates of the human JNK1 isoforms JNK1*α* (369 res., containing exon *6*), JNK1*β* (369 res., containing exon *7*) and JNK1*δ* (304 res., containing neither exon *6* nor exon *7*) were predicted by PhyloSofS pipeline. The 3 systems were prepared with the LEAP module of AMBER 12 [70], using the ff12SB forcefield parameter set: (*i*) hydrogen atoms were added, (*ii*) the protein was hydrated with a cuboid box of explicit TIP3P water molecules with a buffering distance up to 10Å, (*iii*) Na^+^ and Cl^*−*^ counterions were added to neutralize the protein.

The systems were minimized, thermalized and equilibrated using the SANDER module of AMBER 12. The following minimization procedure was applied: (*i*) 10,000 steps of minimization of the water molecules keeping protein atoms fixed, (*ii*) 10,000 steps of minimization keeping only protein backbone fixed to allow protein side chains to relax, (*iii*) 10,000 steps of minimization without any constraint on the system. Heating of the system to the target temperature of 310 K was performed at constant volume using the Berendsen thermostat [71] and while restraining the solute *C_α_* atoms with a force constant of 10 *kcal/mol/*Å^2^. Thereafter, the system was equilibrated for 100 *ps* at constant volume (NVT) and for further 100 *ps* using a Langevin piston (NPT) [72] to maintain the pressure. Finally the restraints were removed and the system was equilibrated for a final 100 *ps* run.

Each system was simulated during 250 ns (5 replicates of 50 ns, starting from different initial velocities) in the NPT ensemble using the PMEMD module of AMBER 12. The temperature was kept at 310 K and pressure at 1 bar using the Langevin piston coupling algorithm. The SHAKE algorithm was used to freeze bonds involving hydrogen atoms, allowing for an integration time step of 2.0 fs. The Particle Mesh Ewald (PME) method [73] was employed to treat long-range electrostatics. The coordinates of the system were written every ps.

Standard analyses of the MD trajectories were performed with the *ptraj* module of AMBER 12. The calculation of the root mean square deviation (RMSD) over all atoms indicated that it took between 5 and 20 ns for the systems to relax. Consequently, the last 30 ns of each replicate were retained for further analysis, totaling 150 000 snapshots for each system. The fluctuations of the C-*α* atoms were recorded along each replicate. For each residue or each system, we report the value averaged over the 5 replicates and the standard deviation (see Fig. 4a). The secondary structures were assigned by DSSP algorithm over the whole conformational ensembles. For each residue, the most frequent secondary structure type was retained (see Fig. 4a and **S7 Fig**). If no secondary structure was present in more than 50% of the MD conformations, then the residue was assigned to a loop. The amplitude of the motion of the A-loop compared to the rest of the protein was estimated by computing the angle between the geometric center of residues 189-192, residue 205 and either residue 211 in the isoforms JNK1*α* and JNK1*β* or residue 209 in the isoform JNK1*δ*. Only C-*α* atoms were considered.

### 1.6. RNA-Seq Data integration

To obtain additional support for transcript isoform expression, we queried the Bgee database (v.14) [74] for a list of all RNA-Seq experiments related to the selected species. Using SRA tools, we downloaded raw sequences from *H. sapiens* (224 samples), *M. musculus* (155 samples), *Xenopus tropicalis* (69 samples) and *D. rerio* (67 samples) and then aligned the reads using STAR v.2.5.3a [75] with default parameters. *T. rubripes* is not annotated in Bgee and was not integrated in this part of the analysis. Reads overlapping exon-exon boundaries (*e.g.* splice-junction reads) next to alternative splicing events provide direct evidence for the expression of specific transcripts isoforms. Combined with sample annotation, they could also inform on tissue specific isoform expression. We thus considered all reads included within one of the JNK genes and monitored the alignment of splice junctions between different exons as support for the transcripts isoforms: exons 5-6 and 6-8 for JN1*α*, exons 5-7 and 7-8 for JNK1*β*, and exons 5-8’ for JNK1*δ*.

## Acknowledgments

We thank Y. Christinat for providing information on the algorithm he developed for the reconstruction of transcript phylogenies.

## Supporting Information Captions

*S1 Text*

*S1 Table*

**3D structures of human JNKs deposited in the Protein Data Bank.**

*S2 Table*

**Overlap between exons and known regions in human JNK tertiary structure.** For each exon, the corresponding residue range in human JNK tertiary structure is indicated, along with the known region(s) overlapping with the exon and the residues from this(ese) region(s) being included in the exon. Exons for which no residue range is indicated (-) map to disordered parts. The horizontal lines separate actual exons (exons *1* and *8* were splitted in *1*, *1’* and *8*, *8’* to model the transcripts in PhyloSofS, see *Methods*).

*S1 Fig*

**Workflow of the transcripts’ phylogeny reconstruction algorithm. (a)** A binary tree representing the phylogeny of the gene(s) of interest is given as input, along with the transcripts observed at the leaves (symbols). Each transcript is described as a collection of exons, each exon being colored differently (white means that the exon is absent from the transcript). **(b)** The first step consists in determining the states of the exons at the level of the gene, either absent (white square), alternative (black/white square) or constitutive (black square). To determine the exon states at the internal nodes, Sankoff’s algorithm and Dollo’s parsimony principle are used. **(c-d)** The algorithm then proceeds iteratively by searching the space of possible forest structures (c) and evaluating the phylogeny of minimum cost for each chosen structure (d). **(c)** A forest structure *S*_*i*_ is fixed by setting the number of binary (with two children), left (with one left child) and right (with one right child) subnodes at each internal node. **(d)** The phylogeny *φ*_*i*_ associated to the forest *S*_*i*_ is computed only if the cost associated to *S*_*i*_, which depends on the number of transcript births and deaths, is lower than the cost *C*_*min*_ of the best phylogeny found so far. At this stage, each transcript is represented by a table of costs, where each line corresponds to an exon and each column corresponds to an exon state. There are four possible states: absent (white square), alternative absent (grey/white square), alternative present (black/grey square) and present (black square). Only the cells permitted by the exon states at the gene level (determined in a) are considered. Sankoff’s algorithm is used bottom up to compute the minimal pairing costs (see Table II for the list of elementary mutation costs). At each internal node, the problem of pairing the children transcripts is that of a partial assignment and is solved by using a branch-and-bound algorithm (see inserted table on the left: the chosen pairs are those with minimum costs and compatible, and *Supplementary text S1*). The total cost associated to mutations along the branches is obtained by summing the costs over all tables, where the cost of each table is the sum of the minimum costs determined for each line (exon). The cost associated to each observed transcript (leaf) is obviously zero.

*S2 Fig*

**Gene tree, species tree and parented human isoforms. (a)** Gene tree for the JNK family, comprising JNK1 (in black), JNK2 (in grey) and JNK3 (in light grey), in 7 species: human, mouse, *Xenopus tropicalis*, fugu, zebrafish, *Drosophila melanogaster* and nematode. Each node represents a gene from a current or ancestral species. The internal nodes are numbered and the leaves are labelled. The fishes contain two paralogs of JNK1 each, named JNK1 and JNK1a in fugu, JNK1a and JNK1b in zebrafish. *Drosophila melanogaster* and nematode contain only one JNK gene and are used as outgroups. **(b)** Mapping of the duplication, loss and speciation events for the JNK family onto the phylogeny of the 7 studied species. The genes being duplicated or lost are indicated on the corresponding branches. **(c)** On top, a simplified representation of the JNK genes structure is displayed. Below, the human isoforms for which a phylogeny could be reconstructed (see Fig. 1b) are listed and the gene(s) producing each isoform is(are) indicated on the left. Rectangles represent exons and are labelled from 0 to 13. Exons colored in black in the gene structure (on top) are present in all the listed isoforms (below) and those in gray are present in only a subset of the isoforms. The splicing paths corresponding to the listed isoforms are highlighted in colors.

*S3 Fig*

**Comparison of exon mapping onto JNK tertiary structure between human and** *Drosophila melanogaster*. The structures of JNK1 from human (**a**, PDB code: 3ELJ [39]) and JNK from *Drosophila melanogaster* (**b**, PDB code: 5AWM [77]) are represented as cartoons. The different exons are colored from blue through white to red. The residues in yellow are at the junction of 2 exons. The exon numbers next to the color strip correspond to those used in PhyloSofS (see *Methods*).

*S4 Fig*

**Multiple sequence alignments of the exons *6* and *7*.** The colors indicate the physico-chemical properties of the amino acids: hydrophobic (AFILMPVW) in red, negatively charged (DE) in blue, positively charged (KR) in magenta, polar or special (CGHNQSTY) in green. The symbols at the bottom of each alignment give the conservation degree of the column: ★ for completely conserved,: for conserved physico-chemical properties,. for variable, and void for highly variable. The numbering on top corresponds to JNK1 and JNK2. The arrows indicate anchor residues found in all kinases.

*S5 Fig*

**Catalytic and regulatory spines. (a)** The PKA kinase (PDB code: 2CPK [78]) is taken as a reference to illustrate the spines identified in all kinases in [42]. **(b-d)** 3D models for the human isoforms JNK1*α* (b), JNK1*β* (c) and JNK1*δ* (d). The residues comprising the spines are displayed as sticks and transparent surfaces and are labelled. The catalytic spine is in yellow. It is anchored by two hydrophobic residues located in the F-helix (in all panels, except for d, JNK1*δ*). The regulatory spine is in red. It is anchored by the N-terminal aspartate of the F-helix, namely D220 in PKA (a) and D207 in JNK (b-c) that forms an H-bond with Y164 (a) or H149 (b-c). In JNK1*δ*, the aspartate is replaced by D282 and the H-bond with H149 is maintained (d).

*S6 Fig*

**Hydrogen-bond pattern associated to the HRD motif backbone strain. (a)** The CDK-substrate complex (PDB code: 1QMZ [79]) is taken as a reference to illustrate the hidden strain identified in kinases structures in [43]. **(b-d)** 3D models for the human isoforms JNK1*α* (b), JNK1*β* (c) and JNK1*δ* (d). Hydrogen-bonds are formed between 3 components: the aspartate in the F-helix, the HRD motif in the catalytic loop and the DFG motif in the activation loop. The 3 component are conserved across the whole protein kinase family.

*S7 Fig*

**Secondary structures for the human JNK1 isoforms.** The secondary structures were recorded along MD simulations of human JNK1*α* (on top), JNK1*β* (in the middle) and JNK1*δ* (at the bottom). For each residue, the most frequent secondary structure type is depicted. If a residue does not adopt a well-defined secondary structure, either *α*-helix or *β*-sheet, for more than 50% of the MD conformations, then it is assigned to a loop.

*S8 Fig*

**H-bond pattern and backbone strain of the HRD motif. (a)** Persistence of the H-bonds formed between H149 and R150 from the HRD motif in the catalytic loop and D207 from the F-helix and D169 from the DFG motif in the catalytic site, recorded along the 5 replicates of MD simulations for JNK1*α*, JNK1*β* and JNK1*δ*. **(b)** Ramachandran plot showing the torsion angle values (phi, psi) of R150 from the HRD motif in the MD conformations of the 3 isoforms.

*S9 Fig*

**Transcripts’ phylogeny reconstructed by PhyloSofS for the JNK family.** The forest is comprised of 7 phylogenetic trees, 19 deaths and 14 orphan leaves. The cost of the phylogeny is 69 (with *C*_*B*_ = 3, *C*_*D*_ = 0 and *σ* = 2). The legend is the same as in Fig. 1b.

*S10 Fig*

**Transcripts’ phylogeny reconstructed by PhyloSofS for the JNK family.** The forest is comprised of 7 phylogenetic trees, 21 deaths and 14 orphan leaves. The cost of the phylogeny is 69 (with *C*_*B*_ = 3, *C*_*D*_ = 0 and *σ* = 2). The legend is the same as in Fig. 1b. Compared to Figure 1b, there is one additional subnode (in orange) in the internal nodes A18 and A24 and one JNK1 transcript in fugu is the leaf of the orange tree instead of the yellow one.

*S11 Fig*

**Transcripts’ phylogeny reconstructed by PhyloSofS for the JNK family.** The forest is comprised of 8 phylogenetic trees, 23 deaths and 13 orphan leaves. The cost of the phylogeny is 69 (with *C*_*B*_ = 3, *C*_*D*_ = 0 and *σ* = 2). The legend is the same as in Fig. 1b. Compared to Fig. 1b, there is one additional tree (in blue, exon composition on the right). The murine JNK3 transcript serving as a leaf for this tree was previously an orphan.

*S12 Fig*

**Prediction of intrinsic disorder in JNK isoforms.** The predictions were performed with DISOPRED [67] (a) and with IUPred [68] (b) on the 2 human JNK3 isoforms colored in green and red on Figure 1b, differing by the inclusion/exclusion of exons *12* or *13*. Exon numbers are indicated on top of the plots. The predictions for exon *13* were added on the right of the 2 plots.

*S13 Fig*

**State diagram illustrating the 12 possible elementary operations that can be applied to a forest structure.** Each elementary operation consists in removing and/or adding subnode(s) to a randomly chosen node in the forest. Each subnode corresponds to a transcript and is represented by a round. It is colored according to the node to which it belongs: the chosen node is in black while its left and right children are colored in orange and green respectively. Deaths are displayed as crosses on the branches. For each transition between 2 states, represented by an arrow, the numbers of binary, left and right subnodes being added or removed are indicated in parenthesis.

