## Supplementary File for "Transcripts’ evolutionary history and structural dynamics give mechanistic insights into the functional diversity of the JNK family"

### Supporting Information: Transcripts evolutionary conservation and structural dynamics give insights into the role of alternative splicing for the JNK family.

#### S1 Text

##### Transcripts assignment algorithm

At each internal node $N$, the problem is that of a partial assignment, where $n_b(N)$ transcripts of the left child $N_l$ must be paired with $n_b(N)$ transcripts of the right child $N_r$. Assignment problems can be solved in polynomial time by using combinatorial optimization algorithms such as the Hungarian method [1]. However, owing to the additional contraints of our problem, we developed a specific version of the branch-and-bound algorithm [2] to solve it. Our algorithm takes as input $n_l$ transcripts of node $N_l$ and $n_r$ transcripts of node $N_r$. In a first step, an assignment cost is computed for each pair of transcripts and stored in table of $n_l \times n_r$ cells. Then, we seek to determine the $n_b(N)$ cells with the smallest costs with no overlapping lines or columns, where $n_b(N)$ is the number of binary subnodes of node $N$, parent of nodes $N_l$ and $N_r$. The algorithm proceeds as follows:

- the branching is performed relative to the cell with minimum cost:
  - either we choose the corresponding assignment and we reduce the dimension of the problem by removing one line and one column for the cost table. The constraints imposed by the forest structure may lead to the removal of several lines or columns;
  - or we forbid the corresponding assignment and we seek for the second minimum for the next branching.
- the bounding is performed by computing the sum of the $n_b(N)$ minimum costs, which represents a lower bound for all solutions that remain to be explored. We stop the exploration if this lower bound is greater than the best solution found so far.

For $n_l = n_s = n$, determining the $k = n_b(N)$ cells with the smallest costs requires to read the $n^2$ cells of the table. The solution consisting in choosing the $k$ smallest cells with no overlapping lines or columns is thus found in $n^2 + (n-1)^2 + ... + (n-k+1)^2 \simeq \mathcal{O}(n^3)$ operations for $k = n$. On the other hand, the exploration of the tree of candidate solutions by systematically forbidding every cell requires $n^2 + n^2 - 1 + ... + n^2 - k + 1 \simeq \mathcal{O}(n^4)$ operations for $k = n$. This gives an idea of the size of the tree of candidate solutions. In practice, this algorithm showed good performance.

Table S1: **3D structures of human JNKs deposited in the Protein Data Bank.**

| Gene | Code_Chain |
|---|---|
| JNK1 | 1UKH_A, 1UKI_A, 2G01_A, 2G01_B, 2GMX_A, 2GMX_B, 2H96_A, 2H96_B, 2NO3_A, 2NO3_B, 2XS0_A, 3ELJ_A, 3O17_A, 3O17_B, 3O2M_A, 3O2M_B, 3PZE_A, 3V3V_A, 4AWI_A, 4E73_A,4L7F_A, 4QTD_A, 4UX9_A, 4UX9_B, 4UX9_C, 4UX9_D, 4YR8_E, 4YR8_A, 4YR8_C, 4YR8_F,2XRW_A, 3VUD_A, 3VUG_A, 3VUH_A, 3VUI_A, 3VUK_A, 3VUL_A, 3VUM_A, 4G1W_A, 4HYS_A, 4HYU_A, 4IZY_A |
| JNK2 | 3E7O_A, 3E7O_B, 3NPC_A, 3NPC_B |
| JNK3 | 1JNK_A, 1PMN_A, 1PMU_A, 1PMV_A, 2B1P_A, 2EXC_X, 2O0U_A, 2O2U_A, 2OK1_A, 2P33_A, 2R9S_A, 2R9S_B, 2WAJ_A, 2ZDT_A, 2ZDU_A, 3CGF_A, 3CGO_A, 3DA6_A, 3FI2_A, 3FI3_A, 3FV8_A, 3G90_X, 3G9L_X, 3G9N_A, 3KVX_A, 3OXI_A, 3OY1_A, 3PTG_A, 3RTP_A, 3TTI_A,3TTJ_A, 3V6R_A, 3V6R_B, 3V6S_A, 3V6S_B, 4H36_A, 4H39_A, 4H3B_A, 4H3B_C, 4KKE_A, 4KKG_A, 4KKH_A, 4U79_A, 4W4V_A, 4W4W_A, 4W4X_A, 4W4Y_A, 4WHZ_A, 4X21_A, 4X21_B,4Y46_A, 4Y5H_A, 4Z9L_A |

Table S2: **Overlap between exons and known regions in human JNK tertiary structure.**

| Exon | | Known region | | Binding site | |
|---|---|---|---|---|---|
| **name** | **residues** | **name** | **residues** | **name** | **residues** |
| *0* | - | | | | |
| *1'*<br>*1* | -<br>1-40 | N-term hairpin<br>P-loop | 10-23<br>35-8 | catal. site | 32-8,40 |
| *2* | 42-84 | C-helix | 63-80 | catal. site | 53,55 |
| *3* | 85-103 | | | catal. site | 86 |
| *4* | 105-150 | | | catal. site<br>D-site | 108-114,117<br>112-3,117-8,121,123,126-7,130-1,133 |
| *5* | 151-205 | A-loop | 169-195 | catal. site<br>D-site<br>F-site | 154-8,168<br>159-163<br>187,189,196-9 |
| *6* | 207-229 | F-helix | 207-220 | F-site | 228-9 |
| *7* | 207-229 | F-helix | 207-220 | F-site | 228-9 |
| *8*<br>*8'* | 231-282<br>283-290 | MAPK insert | 245-263 | F-site | 232,234,253,255-6,258-260,262-3 |
| *9* | 292-332 | | | D-site | 323-4,326,329 |
| *10* | 333-353 | C-term helix | 348-353 | | |
| *11* | 355-379 | C-term helix | 355-362 | | |
| *12* | - | | | | |
| *13* | - | | | | |

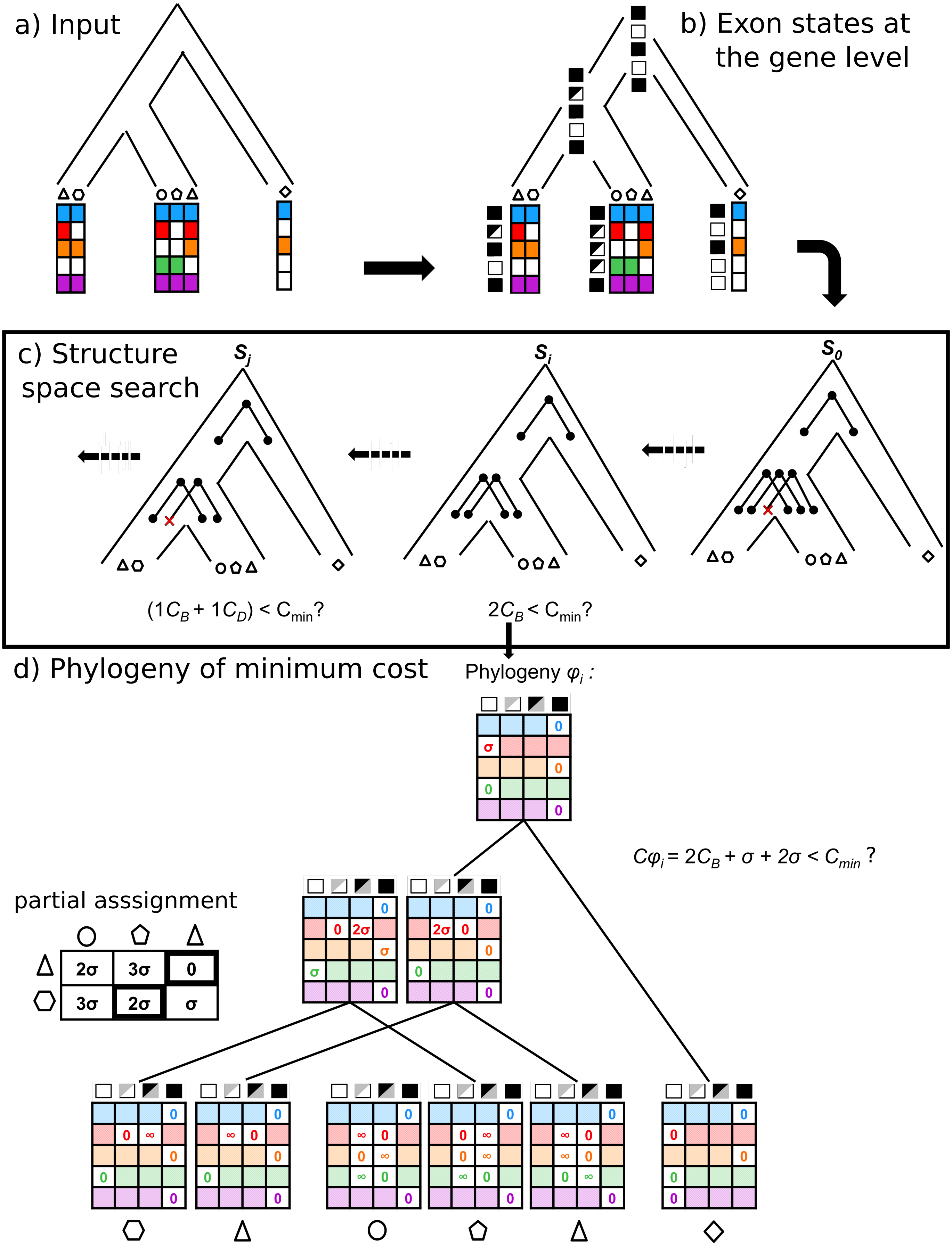

a) Input
b) Exon states at the gene level
c) Structure space search
$S_j$
$S_i$
$S_0$
$(1C_B + 1C_D) < C_{min}$?
$2C_B < C_{min}$?
d) Phylogeny of minimum cost
Phylogeny $\varphi_i$ :
$C\varphi_i = 2C_B + \sigma + 2\sigma < C_{min}$ ?
partial asssignment
2σ
3σ
0
3σ
2σ
σ


Figure S1:

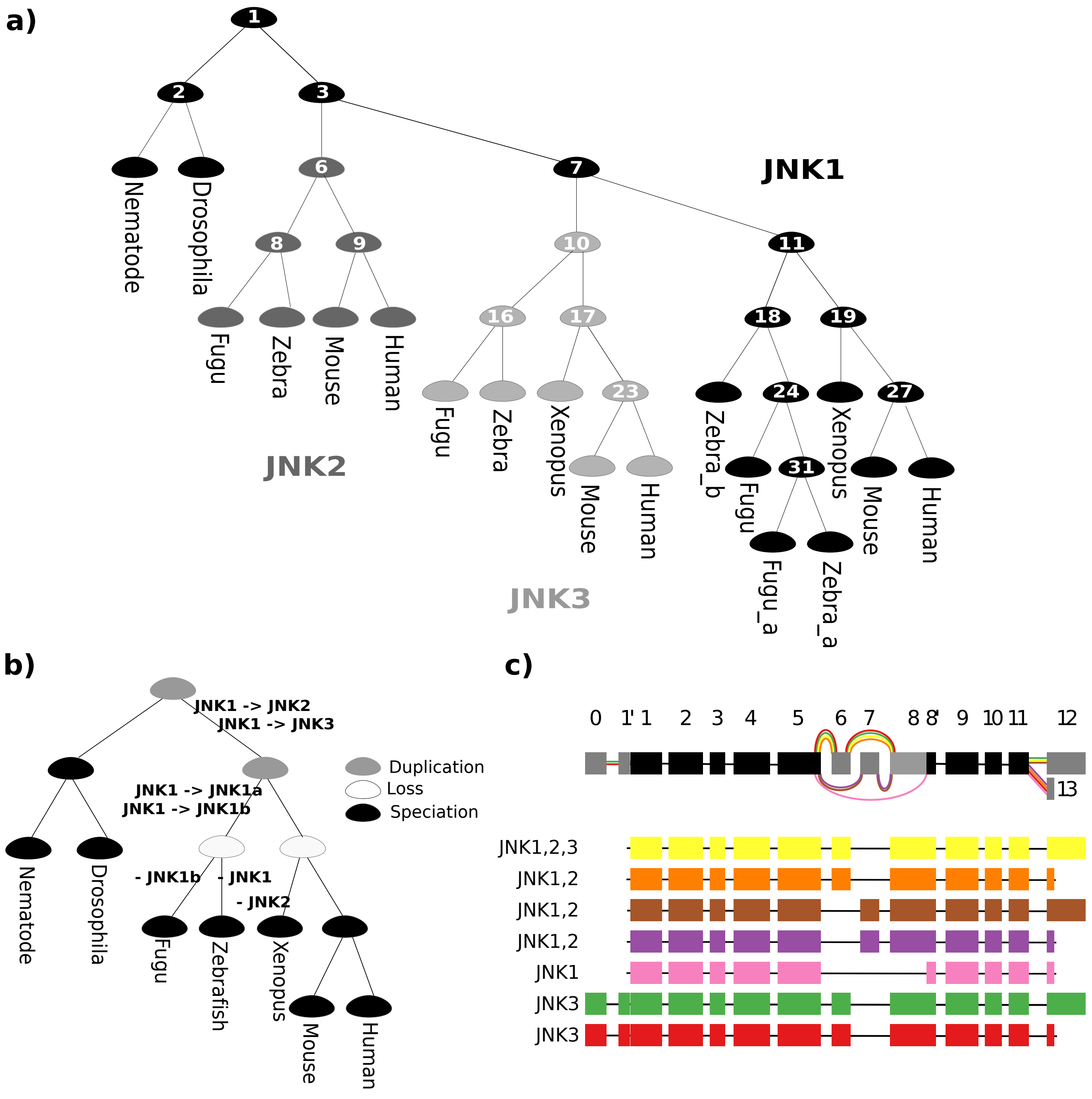


Figure S2: **Gene tree, species tree and parented human isoforms.**

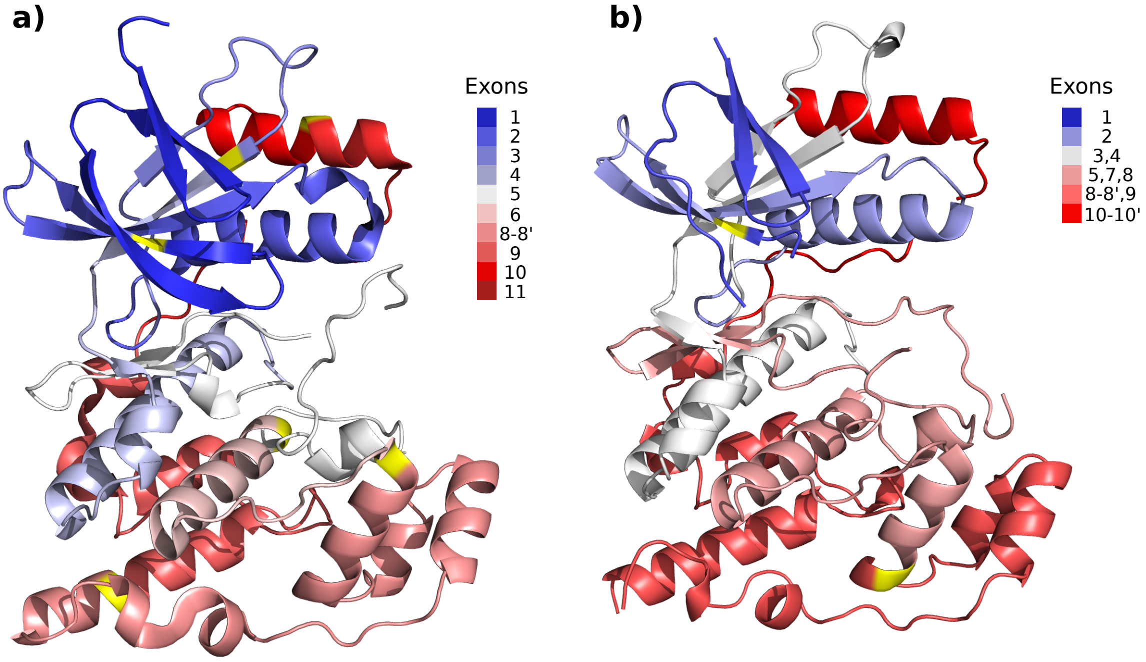


Figure S3: **Comparison of exon mapping onto JNK tertiary structure between human and drosophila.**

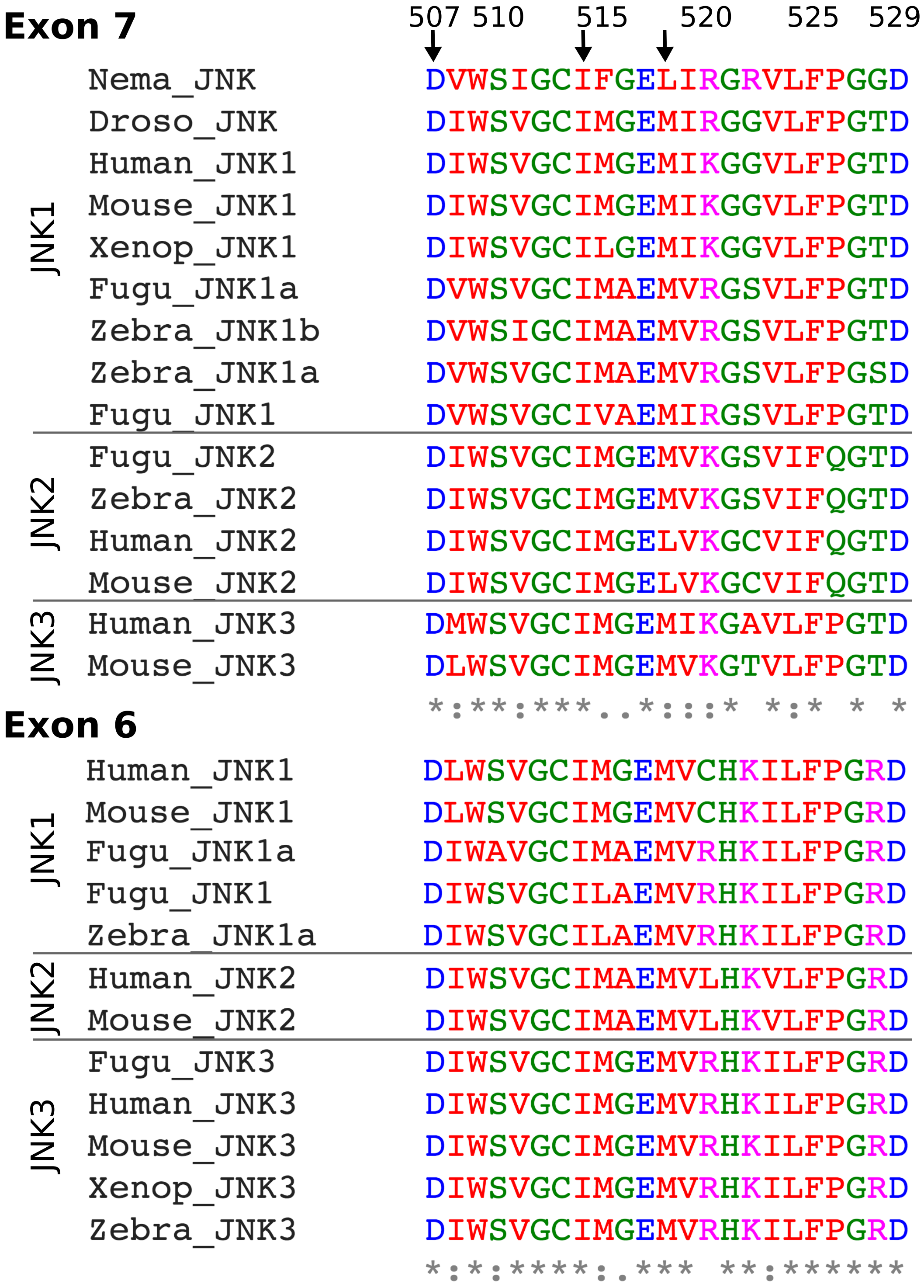


Figure S4: **Multiple sequence alignments of the exons *6* and *7*.**

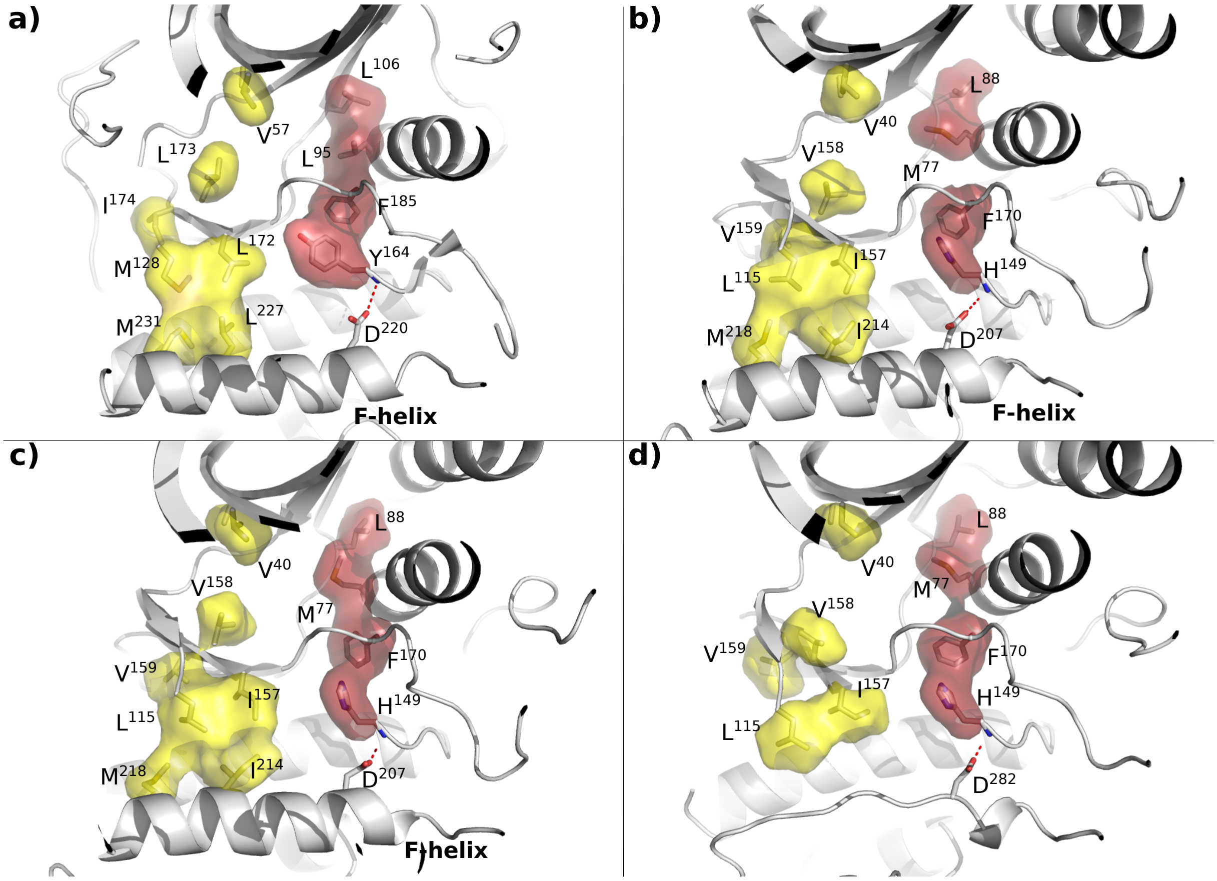


Figure S5: **Catalytic and regulatory spines.**

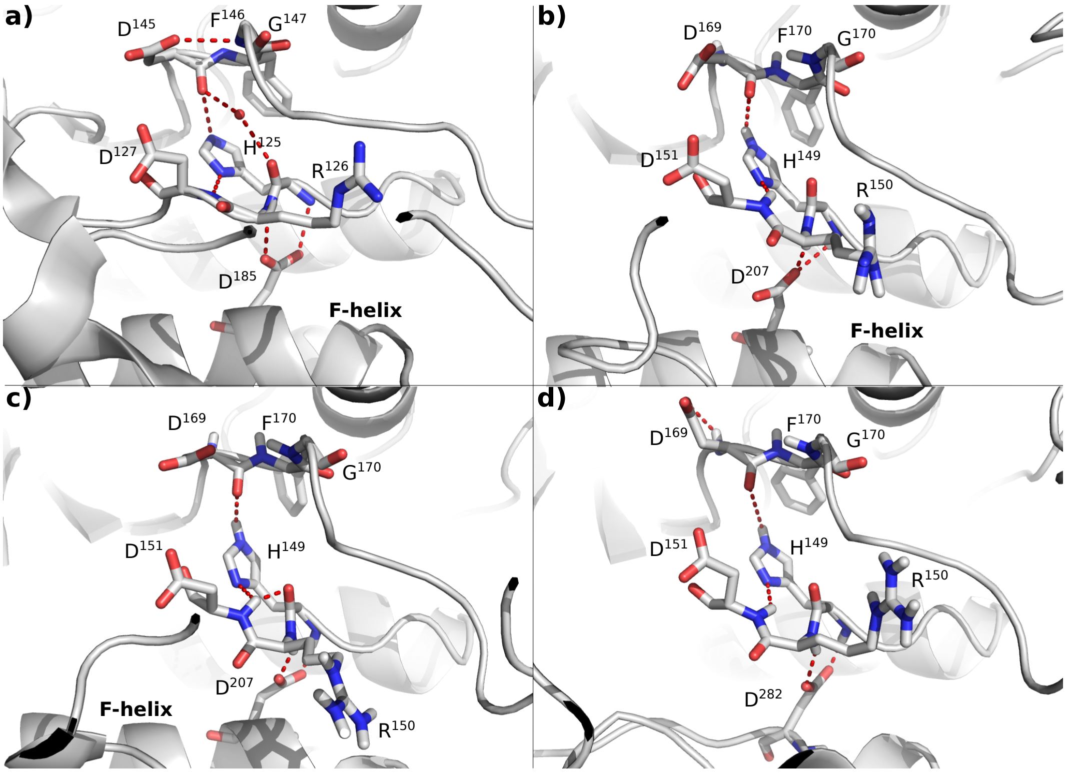


Figure S6: **Hydrogen-bond pattern associated to the HRD motif backbone strain.**

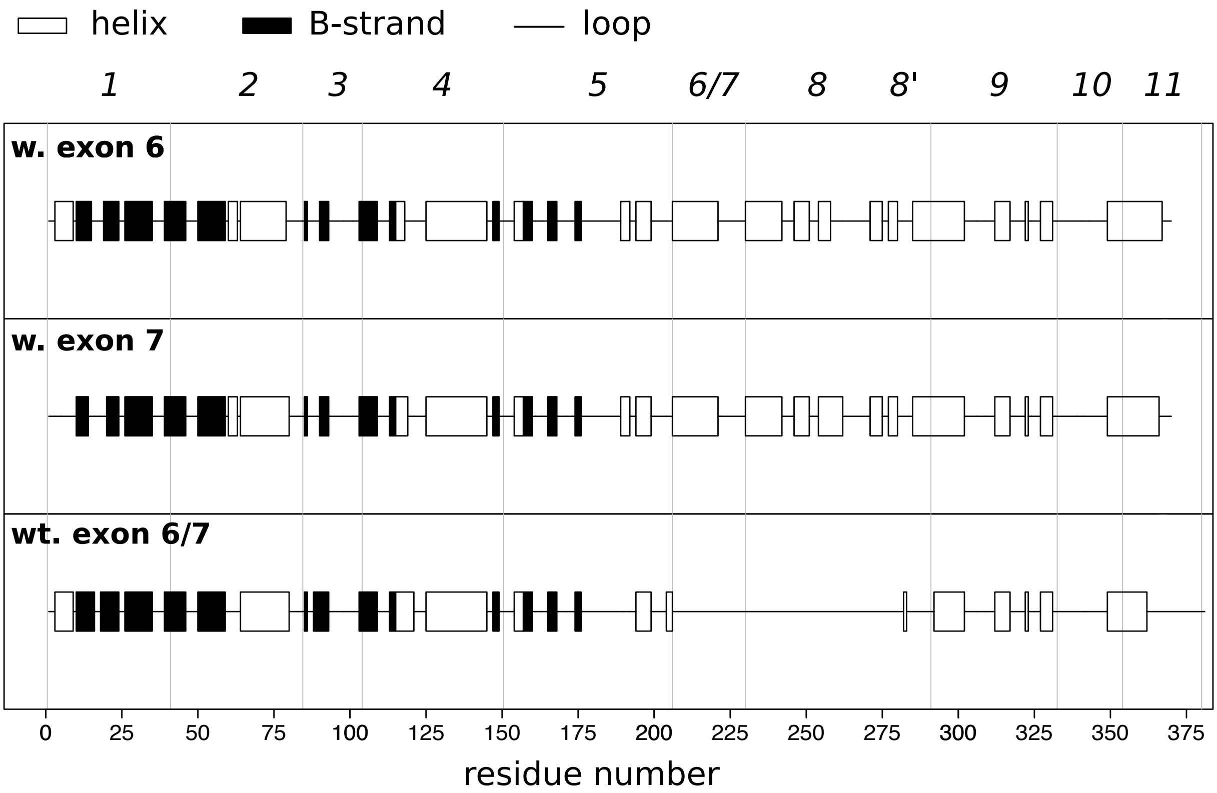


Figure S7: **Secondary structures for the human JNK1 isoforms.**

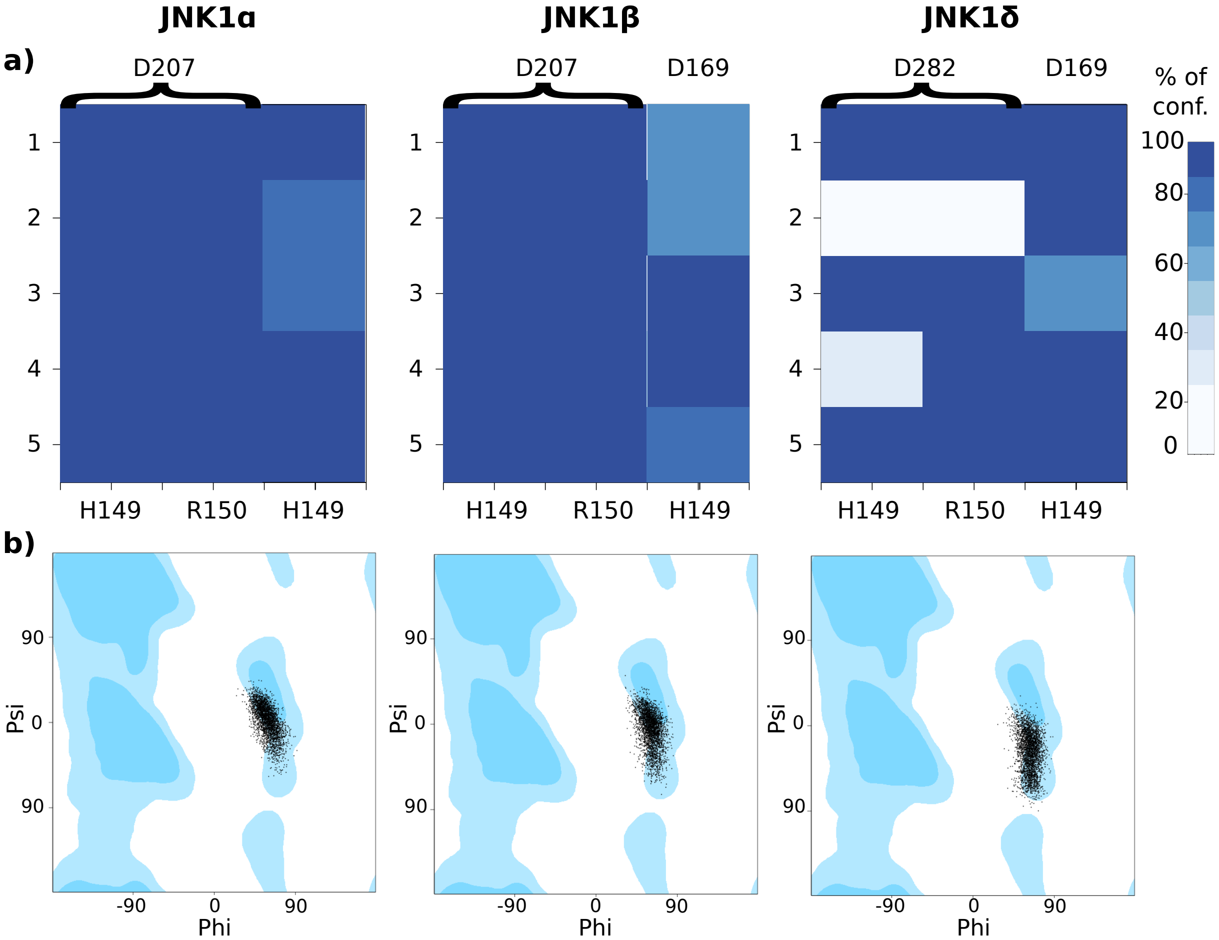


Figure S8: **H-bond pattern and backbone strain of the HRD motif.**

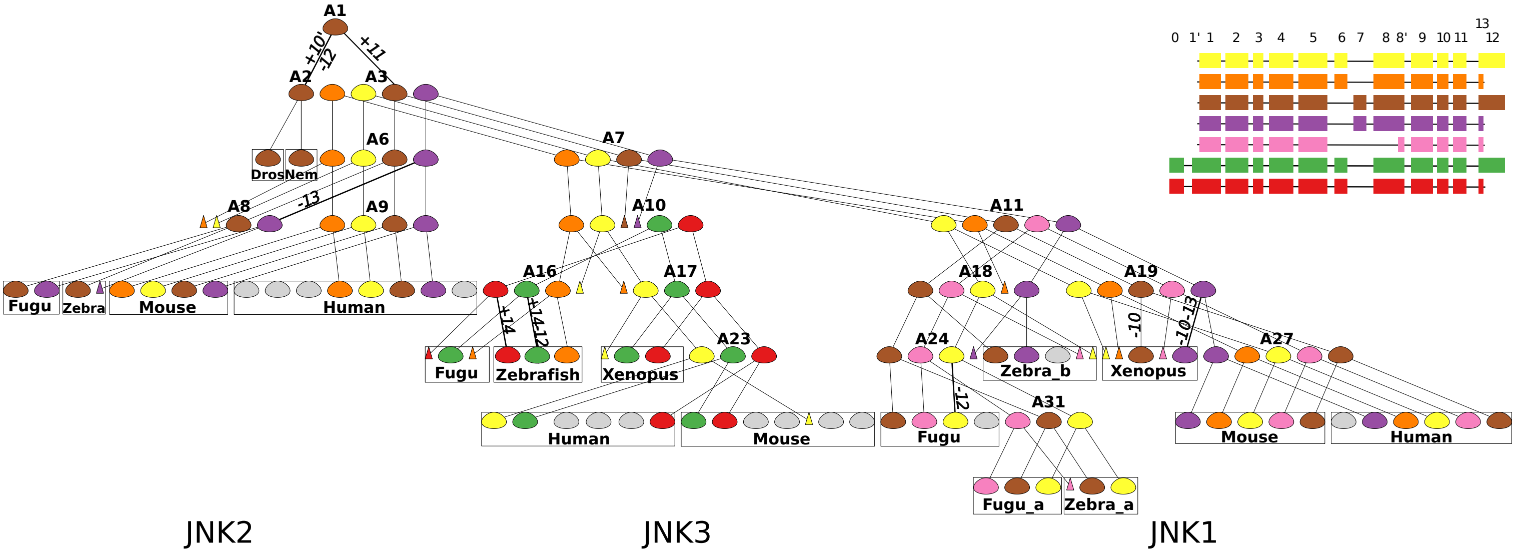


Figure S9: **Transcripts' phylogeny reconstructed by PhyloSofS for the JNK family.**

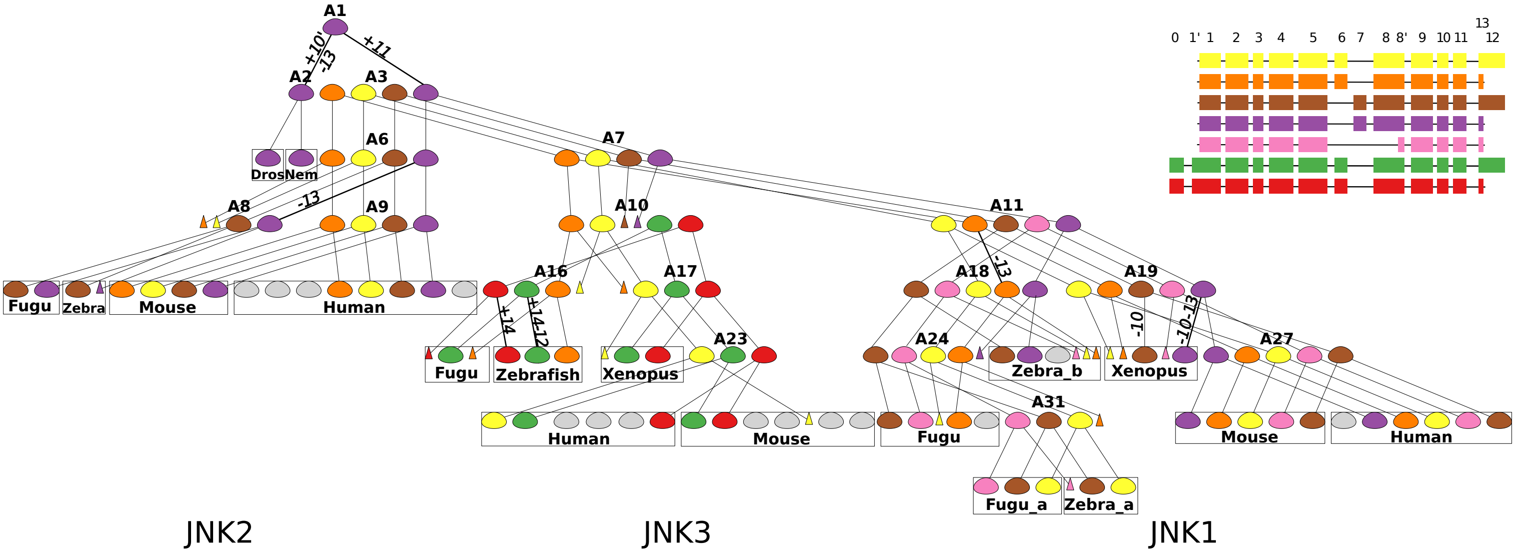


Figure S10: **Transcripts' phylogeny reconstructed by PhyloSofS for the JNK family.**

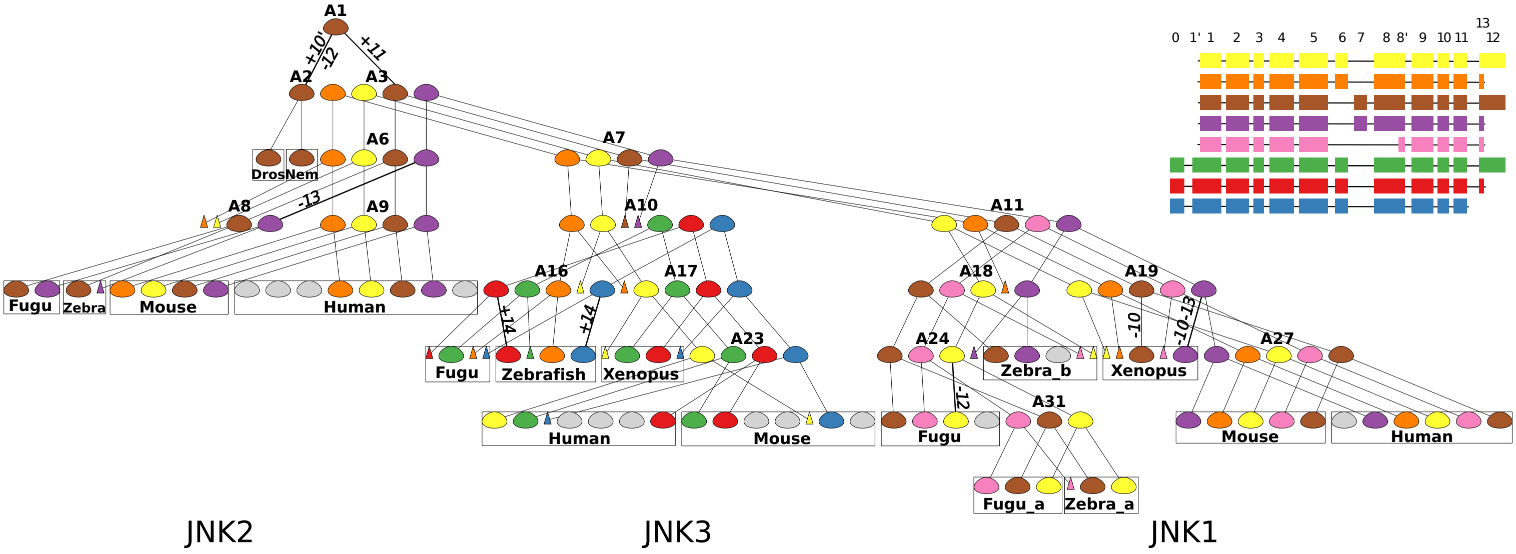


Figure S11: **Transcripts' phylogeny reconstructed by PhyloSofS for the JNK family.**

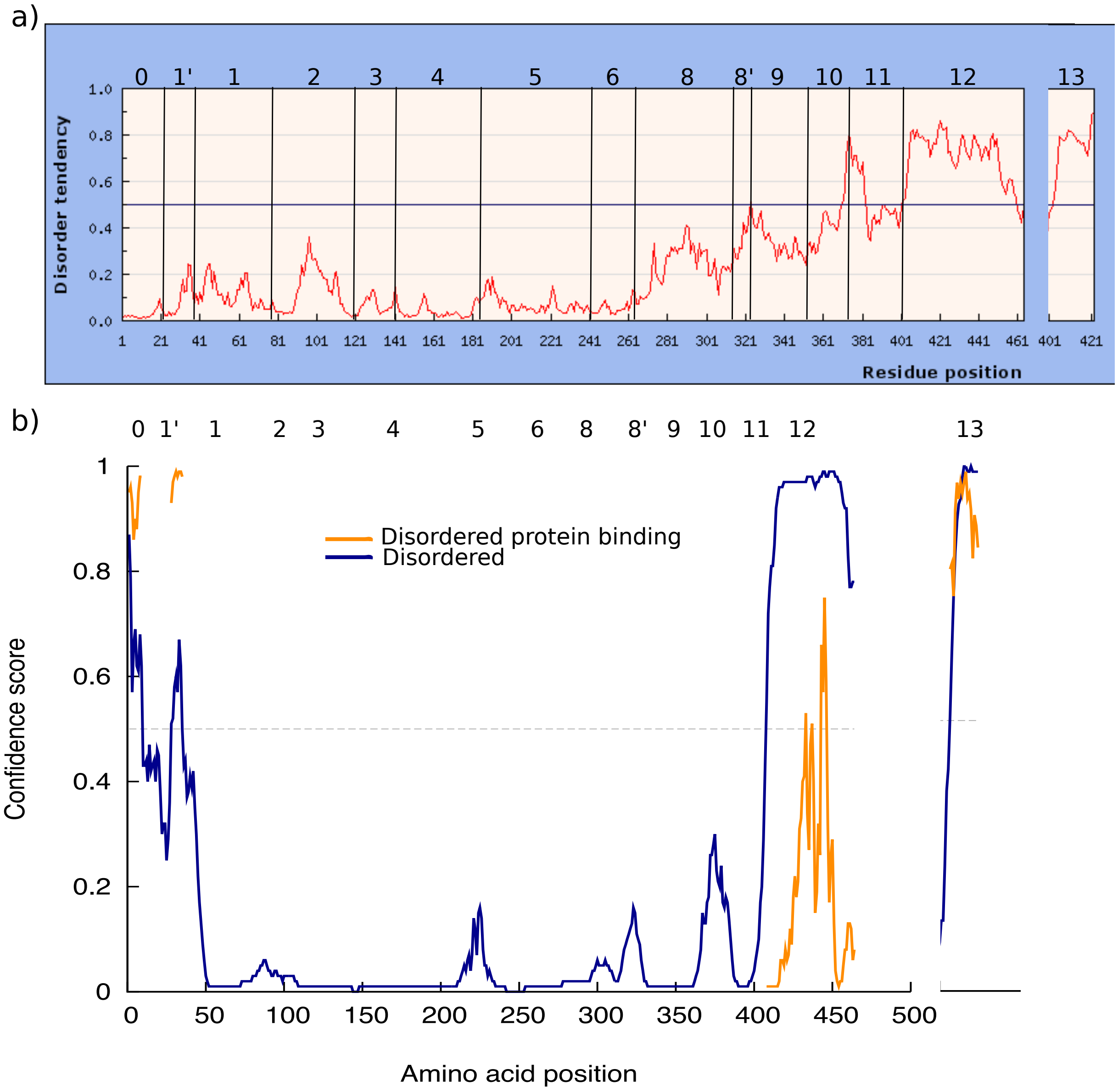


Figure S12: **Prediction of intrinsic disorder in JNK isoforms.**

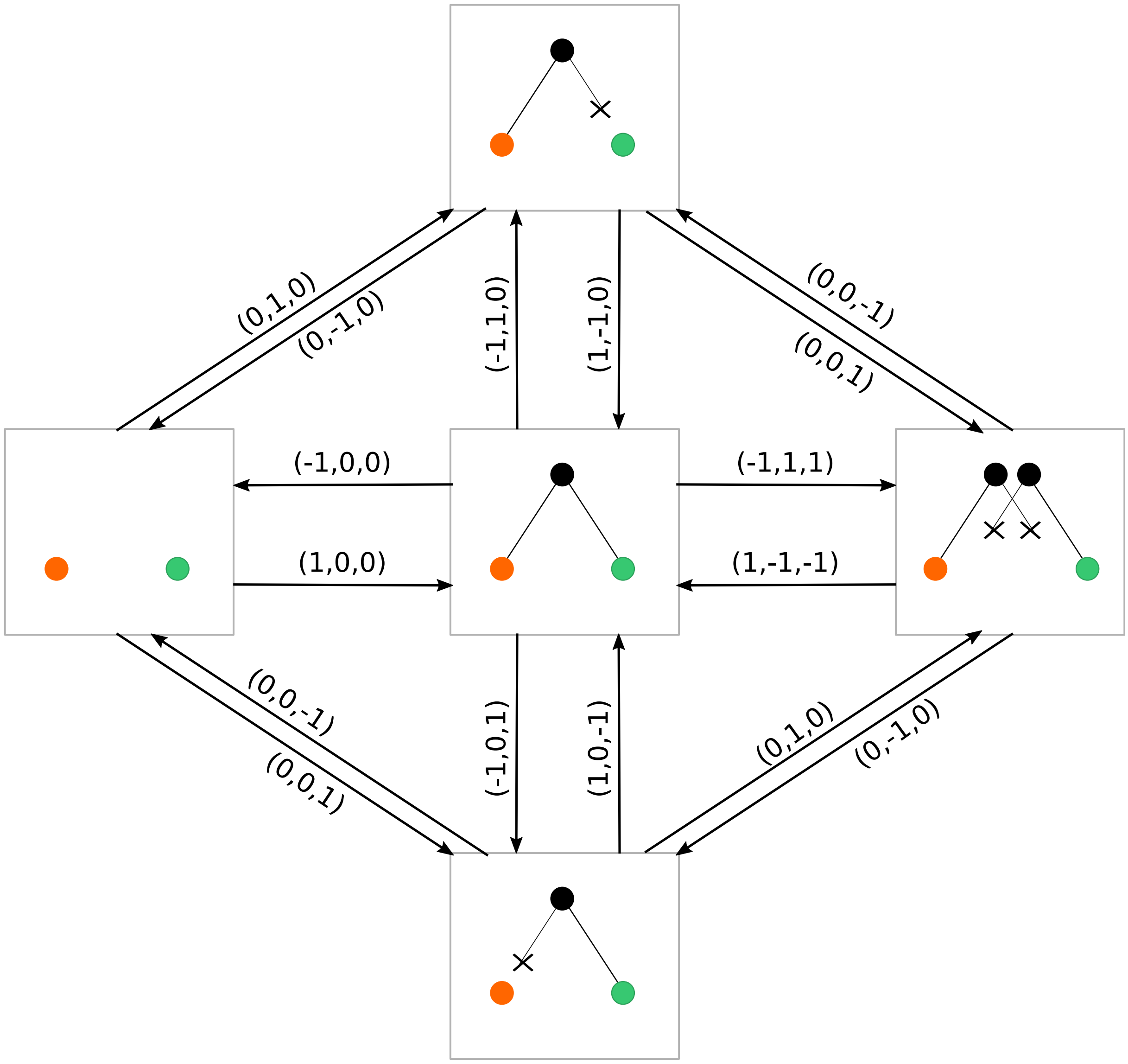


Figure S13: **State diagram illustrating the 12 possible elementary operations that can be applied to a forest structure.**
